# Current progress in understanding Schizophrenia using genomics and pluripotent stem cells: A Meta-analytical overview

**DOI:** 10.1101/2022.08.18.504397

**Authors:** Ashwani Choudhary, Ritu Nayak, David Peles, Liron Mizrahi, Shani Stern

## Abstract

Schizophrenia (SCZ) is a highly heritable, polygenic neuropsychiatric disease, which disables the patients as well as decreases their life expectancy and quality of life. Common and Rare variants studies on SCZ subjects have provided more than 100 genomic loci that hold importance in the context of SCZ pathophysiology. Transcriptomic studies from clinical samples have informed about the differentially expressed genes (DEGs) and non-coding RNAs in SCZ patients. Despite these advancements, no causative genes for SCZ were found and hence SCZ is difficult to recapitulate in animal models. In the last decade, induced Pluripotent Stem Cells (iPSCs)-based models have helped in understanding the neural phenotypes of SCZ by studying patient iPSC-derived 2D neuronal cultures and 3D brain organoids. Here, we have aimed to provide a simplistic overview of the current progress and advancements after synthesizing the enormous literature on SCZ genetics and SCZ iPSC-based models. Although further understanding of SCZ genetics and mechanisms using these technological advancements is required, the recent approaches have allowed to delineate important cellular mechanisms and biological pathways affected in SCZ.

## 1. INTRODUCTION

Schizophrenia (SCZ) is a complex neurological disease that manifests mainly in adulthood although reported to have a neurodevelopment origin (Bloom 1993; Fatemi and Folsom 2009; Ahmad et al. 2018; Schultz, North, and Shields 2007). According to the WHO report, globally approximately 24 million people are affected with SCZ with a worldwide prevalence of around 1% (Stilo and Murray 2010; Rhoades, Jackson, and Teng 2019). SCZ patients are more likely to die younger than the general population mainly due to co-morbid conditions like cardiovascular, diabetes and communicable diseases (Wildgust, Hodgson, and Beary 2010). With the rise in global life expectancy and aging populations, psychiatric disorders like SCZ will further increase the socio-economic liability (Lee et al. 2018), particularly in low and middle-income countries. A few reports have also highlighted the worldwide increase in the incidence of mental disorders like anxiety and depression including psychosis with the COVID-19 pandemic (Xie, Xu, and Al-Aly 2022; Brown et al. 2020; Chacko et al. 2020).

SCZ is characterized by positive symptoms like hallucinations and delusions, negative symptoms like social and emotional withdrawal, and cognitive symptoms like confusion and lack of decision-making (Tandon et al. 2013; Kahn et al. 2015; Page et al. 2022). SCZ is clinically diagnosed according to the criteria mentioned in the DSM V manual based on the characteristic symptoms of hallucinations and delusions; personal and work-related relationships; duration of symptoms and a pre-existing psychiatric condition etc. (Tandon et al. 2013; Kahn et al. 2015). SCZ was observed in relatives and families of patients and is believed to be inherited even before the advent of newer technologies like Genome-Wide association studies (GWAS) and Next-Generation sequencing (NGS) (Henriksen, Nordgaard, and Jansson 2017). Meta-analytical studies of monozygotic and dizygotic twins have suggested an 80% heritability factor for SCZ (Cardno and Gottesman 2000; Sullivan, Kendler, and Neale 2003). Although both candidate gene approaches and high-throughput genomics approaches have identified approximately 170 genomic loci associated with SCZ but the majority are not useful for clinical applications (Henriksen, Nordgaard, and Jansson 2017; Lam et al. 2019; Trubetskoy et al. 2022). An increasing number of reports have formidably associated environmental risk factors in the development of SCZ. Environmental risk factors such as maternal infections and stress as well as maternal malnutrition, which leads to pregnancy and birth complications, also support the neuro-developmental hypothesis (Stilo and Murray 2010; Fatemi and Folsom 2009). Other risk factors for SCZ include socioeconomic status, urban living, increased paternal age, migration status, head injury, autoimmune diseases, epilepsy, and drug abuse (Cannon, Jones, and Murray 2002; Malaspina et al. 2001; Lederbogen et al. 2011).

SCZ is primarily managed clinically with first and second-generation antipsychotics that are known to act on the dopamine D2 receptor and are more effective for positive symptoms (Kane and Correll 2010; Tandon, Nasrallah, and Keshavan 2010). Recent reports show that second-generation antipsychotics, despite being thought to have better efficacy and fewer neurological adverse effects, show comparable efficacy to first-generation antipsychotics and equally increase the incidences of metabolic diseases in SCZ patients (De Hert et al. 2011; Nielsen et al. 2015; Leucht et al. 2013). Treatment-resistant schizophrenia (TRS), which accounts for 30% of all SCZ cases, is treated with Clozapine, which is successful in reducing antipsychotic symptoms, especially in TRS patients, even though the mechanism of action of Clozapine is not fully understood (Potkin et al. 2020; Meltzer et al. 2003). The current gap in the treatment of SCZ and other psychiatric diseases (Kohn et al. 2004) is mainly in the development of drugs that could treat the negative and cognitive symptoms effectively with minimal side effects (Kane and Correll 2010).

Neuroimaging techniques have highlighted structural abnormalities in the brain of SCZ patients. Enlargement of lateral ventricles, thinner cortical areas and reduced hippocampal volumes are established changes found in SCZ patients (van Haren et al. 2011; Haijma et al. 2013). Combining neuroimaging tools with the genetic variants data has been a new field of research to further validate and understand the function of the genetic variants in the SCZ patient subgroups (Callicott et al. 2005). Neuroimaging tools have also been used to validate two environmental risk factors – urban life and migration status (Akdeniz et al. 2014).

Animal models have significantly aided in the understanding of brain development, particularly in informing about evolutionarily conserved cellular and molecular pathways (Pinnapureddy et al. 2015). Although transgenic models have helped in unraveling some aspects of monogenic diseases, the complexity and heterogeneity of polygenic diseases are difficult to replicate in such model systems (Kaiser, Zhou, and Feng 2017). Because of the interactions of multiple genetic and environmental risk factors, neuropsychiatric diseases such as SCZ, Bipolar disorder (BD), and Autism spectrum disorders (ASD) are difficult to study entirely in an animal model system (Nestler and Hyman 2010). At the cellular and molecular level, the use of iPSC technology in neuropsychiatric diseases has provided an alternative strategy to understand the complexity of patient-specific cell lines albeit with limitations as reviewed previously (Soliman et al. 2017; De Los Angeles et al. 2021; Nayak et al. 2021).

In this paper, we present a collective update of the findings from SCZ-iPSC model systems and knowledge gained through GWAS & transcriptomic studies on SCZ clinical samples. Although previous reports and reviews have discussed SCZ iPSC models, GWAS and transcriptomic studies in detail, our goal is to present a quantitative meta-analytic overview of the overall prevailing knowledge to understand the current advancements in the field of SCZ disease pathogenesis and progression.

## 2. DISCUSSION

### 2.1. What insights do genomic studies in SCZ provide?

Recent studies have established the existence of interaction between multiple genetic and environmental factors in the SCZ disease epidemiology (Kahn et al. 2015). Nonetheless, the contribution of genetic factors is large as SCZ is among the highly heritable neuropsychiatric disorder (more than BD and ASD) although without definite causative genes (Sullivan, Daly, and O’Donovan 2012; Schizophrenia Working Group of the Psychiatric Genomics 2014). Using classical genetics techniques some genes were found to be associated with SCZ but in very small sample sizes and which do not follow the Mendelian form of inheritance (Farrell et al. 2015; Sullivan, Daly, and O’Donovan 2012; Trifu et al. 2020). Over the last decade, GWAS has revolutionized the field of complex genetic diseases such as SCZ, by massively genotyping large sample sizes of patients to find genotype-phenotype relationships (Schizophrenia Working Group of the Psychiatric Genomics 2014; Uffelmann et al. 2021). GWAS and NGS have helped in capturing genetic variants, which constitute to partially reveal SCZ genetics (Rhoades, Jackson, and Teng 2019).

#### 2.1.1. Common Variants

Psychiatric genomics consortium (PGC) published studies in large sample sizes to provide common gene variants associated with SCZ. Surprisingly, *miR 137*, a micro-RNA that regulates neural development, was shown to have the highest association with SCZ in the analysis of populations with European ancestry in 2011(Schizophrenia Psychiatric Genome-Wide Association Study 2011a). Their subsequent paper in 2014 with samples from both European and Asian ancestry reported SNPs linked to genes associated with glutamatergic neurotransmission and synaptic plasticity (Schizophrenia Working Group of the Psychiatric Genomics 2014). DRD2 gene, which encodes for the dopamine D2 receptor that is also the known SCZ drug target, was also reported to be significantly associated with SCZ (Schizophrenia Working Group of the Psychiatric Genomics 2014). MHC (Major histocompatibility Complex) region, another important genetic variant, was first found to be associated with SCZ by the Stefansson group (Purcell et al. 2009) and was also reported by subsequent studies (Irish Schizophrenia Genomics and the Wellcome Trust Case Control 2012; Ripke et al. 2013). A recent paper by the PGC consortium has found 120 genes to be involved in fundamental processes such as synaptic organization, neuronal differentiation, and neuronal transmission. Among these genes were the glutamate receptor subunit GRIN2A and transcription factor SP4 and other genes that were linked to rare disruptive coding variants in patients with SCZ (Trubetskoy et al. 2022).

To search for consensus genetic variants across different studies, we have performed a meta-analysis of SCZ GWAS as mentioned in the methods section. The majority of the GWAS on SCZ populations to date has been performed in European and East Asian populations (Fig1.a.). A few studies have also been done on African, Latin American, and South Asian populations among others. Our analysis resulted in a list of highly significant genes that were repeatedly reported as significant across different GWAS publications (p-value <0.01, see Methods) (Fig1.b). CACNA1C was found to be reported in most of the studies along with the HLA locus of the MHC region. CACNA1C is a calcium channel known to have a role in various neurodegenerative diseases (Indelicato and Boesch 2021). It has also been found to be associated with BD in GWAS analysis of common variants (Schizophrenia Psychiatric Genome-Wide Association Study 2011a). NOTCH 4, which has a role in inflammatory pathways, was also found to be highly replicated in multiple studies (Harb et al. 2021). Together with the MHC region, it highlights the important role that the immune system plays in SCZ. Some long intergenic non-coding RNAs were also reported by several studies, further reinforcing the role of non-coding RNAs in regulating gene expression as well as mechanisms that have not yet been fully understood (Fig1.b). TRIM 26 is another important gene that is known to regulate interferon-gamma signaling pathways and is associated with neural tube defects (Zhang et al. 2015; Zhao et al. 2021). However, the discovery of common genetic variants in SCZ has only provided insights into a fraction of SCZ genetics and fuelled a debate about the missing heritability in complex diseases (van Dongen and Boomsma 2013; Zuk et al. 2012; Maher 2008). This has also led to the hypothesis of the presence of undiscovered rare variants with high penetrance (Rhoades, Jackson, and Teng 2019).

#### 2.1.2. Rare variants

Over the years, genetic studies in SCZ patients have been able to identify eleven effective rare structural variants in the genomic region, which have been associated with SCZ and reported to increase the risk of SCZ (Rees et al. 2014). These rare structural variants have neither very high penetrance, nor specificity, as they are also associated with other neurodevelopmental disorders. (Rees et al. 2014; Sullivan, Daly, and O’Donovan 2012) (Table 3). Among these rare variants, the copy number variation of 22q11.21 is well studied and increases the SCZ risk by 20 times but is also associated with Di-George syndrome, developmental delays, and intellectual disability (Cleynen et al. 2021). Most of these structural variants span a large genomic region and are difficult to study in model systems (Table 3). iPSC models of some of these variations have been studied and discussed below.

In addition to the rare structural variants, other rare variants have been identified recently using exome sequencing studies and Whole-genome sequencing approaches. RELN mutations were found to be associated with SCZ in Whole-genome sequencing of a Chinese family with SCZ-affected members. SETD1A loss of function gene variant was found to be significantly associated with SCZ and other developmental disorders in an exome study (Singh et al. 2016). A recent large study on exome sequencing by SCHEMA has found 10 ultra-rare variants in 10 genes, which included representative populations from all seven continents (Singh et al. 2022). Hence, more research using exome sequencing and NGS is required for the exploration of further important genes. Despite these findings, the major portion of SCZ genetics remains undiscovered (Singh et al. 2022). Moreover, the discovered genetic variants have not been helpful in clinical management although recent development of Polygenic risk scores (PRS) profiling for estimating the cumulative effect of small effect-common genetic variants along with rare variants could be helpful in the prediction and intervention of SCZ in clinics in the future (Iyegbe et al. 2014).

#### 2.1.3. Gene expression studies

The lack of discovery of causal genes by GWAS in SCZ makes the study of gene expression pertinent (Rhoades, Jackson, and Teng 2019; Stern, Zhang, et al. 2022). Technological advancements in RNA sequencing have replaced the incomprehensive microarrays-based gene expression assays and allowed for the study of the whole transcriptome of the cells and tissues. A meta-analytical review (Merikangas et al. 2022) has recently reported the list of shortlisted differentially expressed genes in SCZ after analyzing the transcriptomic studies performed from SCZ samples to date. These studies were performed mostly on post-mortem brain tissues and blood of SCZ patients (Fig.1.c). A few studies were also done on LCLs, skin fibroblasts, and iPSCs from SCZ clinical samples (Fig.1.c). The differentially expressed genes were shortlisted by Merikangas et al. (Merikangas et al. 2022) based on their three or more reports of the genes in different studies. We analyzed to find the overlapping shared genes between the highly reported genetic variants as determined by us in Fig 1.b. and the list of 160 differentially expressed genes (DEGs) in SCZ as mentioned by Merikangas et al (Merikangas et al. 2022). We found a list of 14 genes that were common between the highly reported common variant genes and the list of DEGs (Fig1.d). Further analysis of such genes could help in understanding their biological role in SCZ pathophysiology.

Noncoding RNAs have emerged as important biological regulators in brain development and disorders (Qureshi and Mehler 2012) (Salta and De Strooper 2012). As mentioned earlier, SNP associated with miR-137 has been shown to have a strong statistical association with SCZ in GWAS (Schizophrenia Psychiatric Genome-Wide Association Study 2011b) (Fig1.b). In our analysis, we found miR-137 to be differentially expressed in PBMCs and blood of SCZ patients (Fig1.e). Furthermore, miR-7, miR-9, miR-26b, miR-34a, miR-132, miR-181b, and miR-212 were observed to be differentially expressed in a variety of tissues, including PBMCs, blood, and postmortem-brain samples from SCZ patients (Fig1.e). Some iPSC studies have also suggested the role of miRNAs in SCZ iPSC-derived NPCs and Neurons (Fig 1. e). GWAS datasets have also shown an association with long intergenic non-coding RNAs in several studies (Fig1.b). Recently, several lncRNAs have been reported to be differentially expressed in the SCZ clinical samples (Barry et al. 2014; Tian et al. 2018; Liu et al. 2018). Some of the lncRNAs reported in multiple publications have also been listed in Fig 1e. Circular RNAs, a new class of non-coding RNAs, also were shown to be involved in SCZ (Mahmoudi et al. 2019) but not enough studies have been published for meta-analysis according to our criteria (see Methods).

### 2.2. iPSC-based models and how they have helped in understanding SCZ pathophysiology

Since its discovery in the previous decade (Takahashi and Yamanaka 2006), iPSC technology has been utilized to model a variety of diseases, including neurodegenerative and neuropsychiatric disorders(Stern, Sarkar, Galor, et al. 2020; Stern, Sarkar, Stern, et al. 2020; Stern, Zhang, et al. 2022; Brant et al. 2021; Quraishi et al. 2019; Stern, Lau, et al. 2022; Rowe and Daley 2019). It has a considerable advantage in the research of fundamental mechanisms and pathways affecting neuropsychiatric illnesses such as SCZ, BD, and ASD that are otherwise difficult to model due to a lack of causative genes (Soliman et al. 2017; Stern, Sarkar, Stern, et al. 2020; De Los Angeles et al. 2021). Chiang et al. first reported the generation of iPSCs from an SCZ patient harboring a DISC1 mutation (Chiang et al. 2011). We have performed a quantitative meta-analysis for iPSC-based models of SCZ in Fig 2a (i-iii) and Fig 2b. as described in the Methods section by analyzing the findings of more than 50 papers published since the first report in 2011. The relevant phenotypes observed among the different SCZ-iPSC models are plotted in Fig. 2. b. iPSCs can be derived from a variety of somatic tissue but the majority of the research groups have utilized fibroblasts for reprogramming control and SCZ patient samples (Fig.2a.i.). The seminal articles used both lentiviral and episomal vectors to derive iPSCs for SCZ-iPSC models (Chiang et al. 2011; Brennand et al. 2011). Integration-free reprogramming approaches such as Sendai virus vectors and mRNA reprogramming methods have also been successfully employed for reprogramming into iPSCs (Fig.2.a.ii.).

Different techniques related to transcriptomic analysis (RNA-seq, scRNA-seq), gene editing (CRISPR/Cas9, TALENs), high-resolution microscopy, and functional analysis (Multielectrode arrays, Calcium imaging, and Patch-clamp) have been used with iPSCs, neural progenitor cells (NPCs), and neurons derived from SCZ patients to study the mechanisms underlying SCZ pathology. iPSCs are typically differentiated into neural cells through Embryoid Bodies (EBs), neuronal rosettes, and NPCs using specific chemicals and small molecules, and thereafter terminally differentiated into neuronal subtypes using neurotrophic and specific growth factors to obtain neural cell types. For NPC/ Neural Stem Cell (NSC) induction from iPSCs, dual SMAD inhibition has become the preferred method (Chambers et al. 2009). The majority of the studies were conducted on cortical glutamatergic neurons or mixed populations containing both excitatory and inhibitory neurons (Fig.2.a.iii.). A few reports also used overexpression of specific transcription factors to directly induce excitatory or inhibitory neurons from iPSCs to study the SCZ disease mechanisms (Li et al. 2019; Sellgren et al. 2019; Pak et al. 2021). Some studies generated cerebral brain organoids to further understand the SCZ disease pathology in a model, which better mimics brain structure (Fig.2.a.iii) (Kathuria et al. 2020; Stachowiak et al. 2017; Sawada et al. 2020). Here, we briefly discuss the disease relevant findings from SCZ-iPSC-based models, which are also presented in Fig.2.b in the form of a heatmap (described in the Methods section). The heatmap summarizes SCZ phenotypes, which were found to be either upregulated or downregulated among various neural cell types studied in SCZ-iPSC-based model studies.

#### 2.2.1. Disease-relevant phenotypes observed in SCZ iPSC models

##### Abnormal NPCs, Neurites, and Synaptic connections

Brennand et al. conducted the first detailed study using an iPSC-based model to investigate SCZ in 2011, reporting reduced connectivity and neurite extension as well as lower expression of post-synaptic density protein 95 (PSD95) in iPSC-derived cortical neurons from patients with a family history of SCZ (Brennand et al. 2011). Transcriptomic analysis pointed towards significant changes in gene expression related to glutamate, Wnt and cAMP signaling pathways (Brennand et al. 2011). Furthermore, treatment with the antipsychotic drug loxapine improved neuronal connectivity and restored altered gene expression to some extent (Brennand et al. 2011). *In vitro* studies using iPSC-derived neurons and monocyte-derived microglia-like cells from SCZ patients have shown defective synaptic pruning in addition to abnormal synaptogenesis (Sellgren et al. 2019). Synaptic deficits, as well as change in redox state, were also observed in iPSC-derived cortical interneurons of idiopathic SCZ patients (Kathuria et al. 2019a). Treatment with N-acetyl serine, an antioxidant, was found to restore the observed synaptic deficits in SCZ patient-specific cortical interneurons (Kathuria et al. 2019a). In a recent study of discordant monozygotic twin pairs with SCZ, abnormal dendritic arborization resulted in hypoexcitable neurons and synaptic deficits in iPSC-derived dentate gyrus (DG) neurons of co-twin affected with SCZ compared to unaffected co-twin siblings. Moreover, gene expression analysis showed alteration of the DG development pathway, Wnt signaling pathway, and synapse-related pathways in the iPSC-derived DG neurons of SCZ-affected co-twins (Stern, Zhang, et al. 2022).

SCZ-iPSCs-based models with highly penetrant genetic risk factors have shed some light on the cellular phenotypes and genetic pathways. As previously stated, a chromosome 22q11.2 deletion is a known high-risk factor for SCZ. iPSCs derived from the 22q11.2 deletion possessed decreased differentiation ability (Pedrosa et al. 2011; Toyoshima et al. 2016). In addition, smaller neurosphere size and relatively poor neurite extension were also observed after differentiation of the patient iPSCs with 22q11.2 deletion compared to controls (Pedrosa et al. 2011; Toyoshima et al. 2016). In the 15q11.2 deletion iPSC model, another known SCZ risk factor, NPCs lacked apical polarity and adherent junctions indicating a disruption in neuronal development (Yoon et al. 2014). Additionally, iPSC-based cell models were used to study DISC1 mutation as it also carries an increased risk of SCZ (Wen et al. 2014). Synaptic vesicle release was observed to be dysfunctional in iPSC patient-derived neurons with mutant DISC1 compared to controls, and this was rescued when DISC1 was selectively edited in the patient cell line using TALENs, indicating that this mutation affects synaptic activity (Wen et al. 2014). In addition, TALEN and CRISPR-Cas9 editing were employed to introduce the DISC1 mutation into iPSCs resulting in an aberrant neuronal differentiation and a dysregulation of the Wnt signaling pathway as reported earlier by other groups (Srikanth et al. 2015). Exonic deletions of Reelin (RELN) have been linked to SCZ (Costain et al. 2013). In a recent study, glutamatergic and GABAergic iPSC-derived neurons from an SCZ patient with a RELN deletion were reported to have dendritic shortening and fewer synapse counts(Ishii et al. 2019). Those patient-derived neurons were compared to neurons differentiated from isogenic iPSCs specifically edited to have RELN deletion using CRISPR/Cas9 technology (Ishii et al. 2019). The cellular phenotypes of neurons differentiated from genetically edited iPSCs with RELN deletion were comparable with iPSCs of SCZ patients with RELN deletion, confirming the involvement of this gene in SCZ (Ishii et al. 2019). Another study investigating the role of SCZ risk factor, NRXN1 deletion in SCZ-related phenotypes, found altered neurotransmitter release in neurons directly induced from SCZ patient iPSC. They also found neurotransmitter release impairment in induced neurons when iPSCs were edited to have a heterozygous deletion of NRXN1 using TALENs. Importantly, no such defects were observed in mouse neurons deficient of NRXN1 gene reinforcing the importance of human iPSC model for SCZ disease phenotypes (Pak et al. 2021).

##### Mitochondrial dysfunction

An increasing number of reports have linked mitochondrial dysfunction to SCZ (Dror et al. 2002; Prabakaran et al. 2004; Iwamoto, Bundo, and Kato 2004; Rosenfeld et al. 2011). Mitochondrial protein expression and a reduction in membrane potential were reported in SCZ iPSC NPCs, which suggested a compromised mitochondrial function (Robicsek et al. 2013). Altered mitochondrial morphology in SCZ iPSC-NPCs such as smaller and less connected mitochondria and a reduction in mitochondrial density around the nucleus has also been observed (Brennand et al. 2015). Other researchers reported a higher mitochondrial oxygen consumption (Da Silveira Paulsen et al. 2012) and increased reactive oxygen species (ROS) levels (Robicsek et al. 2013; Brennand et al. 2015) in SCZ iPSC-derived NPCs. Robicsek et al. in a subsequent study used isolated active normal mitochondria (IAN-MIT) technology to transfer healthy mitochondria to SCZ iPSCs which rescued the impaired differentiation of SCZ-iPSCs into glutamatergic neurons (Robicsek et al. 2017). A study found a reduction in ATP levels due to reduced oxidative phosphorylation at mitochondrial complexes I and IV in the forebrain excitatory neurons directly induced from iPSCs from 22q11.2 SCZ subjects (Li et al. 2019). Furthermore, mitochondrial protein expression was specifically altered and not the nuclear-encoded protein in those neurons (Li et al. 2019). Ni *et al*. found dysregulation of oxidative phosphorylation genes and altered mitochondrial function in cortical interneurons differentiated from SCZ patient-derived iPSCs compared to iPSCs derived from the control group (Ni et al. 2020). Surprisingly, they observed no such significant changes in glutamatergic neurons derived from the same SCZ patient iPSCs. Furthermore, Alpha Lipoic Acid/Acetyl-L-Carnitine (ALA/ALC) was found to restore the observed dysregulation of oxidative phosphorylation genes in cortical interneurons (Ni et al. 2020). These observations suggest that alterations in Mitochondria structure and function may play an important role in SCZ pathology.

##### Altered Neuronal functions

Evidence for aberrant glutamate and GABA signaling in neurons derived from SCZ iPSC have been provided by several studies. Both transcriptomic and proteomic analyses have found dysregulation in receptor proteins as well as differential expression of glutamate and GABA signaling pathway genes (Brennand et al. 2011; Brennand et al. 2015; Kathuria et al. 2019b; Kathuria et al. 2020; Narla et al. 2017; Tiihonen et al. 2019). Calcium imaging and microelectrode array experiments have been used to measure the altered excitatory and inhibitory responses of SCZ patient-specific neurons in several studies (Tiihonen et al. 2019; Kathuria et al. 2019b; Kathuria et al. 2020; Hathy et al. 2020; Stern, Zhang, et al. 2022; Sarkar et al. 2018). Electrophysiological recordings have found reduced activity in iPSC-derived neurons from SCZ patients as well as changes in excitatory and inhibitory synaptic activity (Fig 2b). Clozapine was found to affect the differential calcium response to glutamate and GABA observed in SCZ iPSC-derived neurons (Tiihonen et al. 2019). Recent studies have found associations between electrophysiological measures and SCZ clinical status in iPSC-based models (Page et al. 2022). Cortical neurons derived from SCZ patients had altered sodium channel function and firing potential and increased GABA transmission as compared to controls (Page et al. 2022).

##### Findings from 3D models of SCZ brain organoid

Cortical malformation was observed in brain organoids of three SCZ patients and linked to FGFR1 signaling (Stachowiak et al. 2017). In addition, expression of the Reelin protein, known to have an important role in corticogenesis and earlier reported to be deficient in SCZ patients and 2D SCZ-based models, was found to be decreased in the SCZ patient-derived organoids (Stachowiak et al. 2017). The brain organoid studies from monozygotic twins with SCZ revealed changes in the excitatory and inhibitory balance with an increase in GABAergic synaptic genes compared to healthy controls (Sawada et al. 2020). NPC proliferation and changes in the ventricles/rosettes per organoid were observed in DISC1 deficient SCZ brain organoids as compared to controls (Srikanth et al. 2018). Also, WNT signaling genes were found to be dysregulated in the transcriptomic analysis of these SCZ-iPSC brain organoids providing mechanistic clues to these observed cellular phenotypes (Srikanth et al. 2018). A recent transcriptomic study of cerebral organoids from SCZ patients found dysregulation of a few SCZ-associated genes that were previously found in related GWAS as well as the alteration in the genes related to mitochondrial function (Kathuria et al. 2020). The mitochondrial functional assay revealed alteration in Mitochondria-oxygen consumption rate in SCZ brain organoids along with changes in excitatory and inhibitory synaptic function measured via microelectrode arrays (Kathuria et al. 2020). Another recent proteomic study using brain organoids derived from SCZ patients found proteins involved in axonal pathways and morphogenesis to be depleted in SCZ. Two proteins (Pleiotrophin and Podocalyxin)reported earlier in GWAS studies were also found to be altered (Notaras et al. 2021). Together, these studies suggest alterations in NPC proliferation/differentiation, alterations of neurites and synaptic connections, and dysregulation of mitochondrial function in SCZ.

##### Insights from Environmental risk factor studies

Prenatal stress is known to be an important environmental risk factor for SCZ. Maternal immune activation (MIA) is a form of prenatal stress that can result from microbe infection during pregnancy (Brown and Derkits 2010; Estes and McAllister 2016). MIA has been associated with an increase in the copy number of LINE-1 retrotransposons in animal models as well as SCZ iPSC-derived neurons from 22q.11 deletion patients (Bundo et al. 2014). LINE-1 retrotransposons are known to create somatic heterogeneity in neurons and were found to be dysregulated in SCZ postmortem brain samples (Doyle et al. 2017). Herpes simplex virus type 1 (HSV-1) infection of SCZ iPSC-derived neurons altered the expression of genes related to glutamatergic signaling and mitochondrial function compared to control iPSC-derived neurons which discloses the aggravating role of viral infection on SCZ genetic background (D’Aiuto et al. 2014). Rodent models have shown induction of several cytokines in the fetal brain after MIA indicating a neuro-inflammatory response (Estes and McAllister 2016). In experiments mimicking neuro-inflammation, IFN gamma treatment to the human iPSC-derived NPCs reflected the similar transcriptomic response reported in SCZ and Autism (Warre-Cornish et al. 2020). Another report studying the effect of activated microglia conditioned media on SCZ-iPSC-derived neurons found differential responses in cortical interneurons and glutamatergic neurons (Park et al. 2020). Cortical interneurons showed changes in gene expression related to metabolic pathways as well as mitochondrial dysfunction, reduced arborization, and synapse deficits when exposed to activated microglia conditioned media while the glutamatergic neurons derived from the SCZ–iPSCs did not change significantly in the study (Park et al. 2020). Moreover, in the presence of mitochondrial function-enhancer compounds (Alpha Lipoic Acid/Acetyl-L-Carnitine), these deficits in cortical interneurons were found to be rescued (Park et al. 2020). But besides these preliminary findings, more exhaustive iPSC-based research into the impact of environmental risk factors on SCZ-relevant biological pathways is required. In addition, the 2D iPSC-based models do not recapitulate the *in vivo* scenario of immunological and inflammatory response that necessitates a coordinated response from neuronal, immunological, and glial cells (Balan, Toyoshima, and Yoshikawa 2019).

#### 2.2.2. Key cellular and molecular pathways identified

As discussed above, several studies have reported phenotypes as well as molecular signatures relevant to SCZ pathology in NPCs and differentiated neural cells from SCZ iPSC-based models (Fig .2.b).) A WNT signaling pathway dysregulation has been consistently reported by various groups employing both 2D and 3D iPSC-derived culture and organoids as well as postmortem clinical samples through transcriptomic and proteomic studies (Sawada et al. 2020; Topol et al. 2015; Brennand et al. 2011; Srikanth et al. 2015; Srikanth et al. 2018; Panaccione et al. 2013). Prior cellular and animal model studies have established the role of WNT signaling in cortical neurogenesis (Wrobel et al. 2007; Mutch et al. 2009). Hence, the dysregulation of the WNT signaling pathway may contribute to the observations of aberrant NPC proliferation/migration and synaptic alterations.

Another consistent observation in many studies has been mitochondrial dysfunction and oxidative stress (Brennand et al. 2015; Kathuria et al. 2020; Robicsek et al. 2013; Da Silveira Paulsen et al. 2012; Ni et al. 2020). Since normal mitochondrial function is important for neuritogenesis and axonogenesis, the alterations of the mitochondrial oxidative pathway may contribute to abnormal neurite and dysregulation of axonal pathways that were observed in SCZ-iPSC-based models (Leon et al. 2016). A few SCZ iPSC studies discussed above have also tested this proof of concept in rescue experiments by enhancing the mitochondrial function using compounds or transferring healthy mitochondria into iPSC-derived neurons (Park et al. 2020; Robicsek et al. 2017; Ni et al. 2020). Moreover, oxidative stress was found to regulate the WNT signaling pathway in neuronal and non-neuronal cells (Zhang, Tannous, and Zheng 2019; Almeida et al. 2007). In addition, oxidative stress is also known to regulate synaptic plasticity (Massaad and Klann 2011) and affect synaptic transmission in Alzheimer’s’ disease (Kamat et al. 2016). N-acetyl serine, an antioxidant, was found to rescue morphological alterations of synapses in SCZ patient-derived cortical interneurons indicating the role that oxidative stress plays in synaptic impairments in SCZ (Kathuria et al. 2019a). We, therefore, speculate that oxidative stress also plays an important role in the observed phenotypes in SCZ. However, these events may be interconnected and influence one another. The contribution of glial cells to SCZ pathology was not studied extensively but in a relatively small number of studies (Akkouh et al. 2020; Akkouh et al. 2021; Windrem et al. 2017; Szabo et al. 2021). Mice implanted with human iPSC-derived glial cells from SCZ patients were observed to have abnormal behavior including anxiety and reduced social interaction (Windrem et al. 2017). The transcriptome analysis of differentiated glial progenitor cells of SCZ patients grown *in vitro* also showed dysregulation of genes related to glial differentiation and synaptic pathways in the same study (Windrem et al. 2017).

Although these lines of evidence point toward the underlying pathophysiology of SCZ, it is important to note that SCZ-based iPSC models have been derived from SCZ patients with different subtypes (Idiopathic or genetic) and have varied clinical phenotypes. Hence, it is important to further stratify the SCZ clinical samples and delineate the important pathways in each clinical subgroup for the designing and discovery of better therapeutic alternatives.

## 3. CONCLUSIONS

Advancements in genomics have increased the understanding of complex polygenic diseases like SCZ. GWAS and transcriptomic studies have significantly helped in the discovery of genetic variants and molecular pathways important in the pathophysiology of SCZ. While the causative genes of SCZ are still obscure, the discovery of common genetic variants from GWAS can be utilized for calculating PRS for further stratifying the SCZ patients’ samples based on their association with clinical symptoms (Zheutlin et al. 2019; Page et al. 2022). Apart from patient-specific cell lines from clinically diagnosed individuals, rare but highly penetrant mutations discovered via genetic studies & implicated in SCZ have been modeled using iPSCs to study the SCZ disease-specific neural cells *in vitro*. Rare genetic variants have aided in understanding the crucial genetic and molecular pathways but some of them are known to have a pleiotropic effect and are also associated with other neuropsychiatric diseases. Genome editing tools are still technically challenging for large genomic deletions and allow currently for the editing of small genomic regions that are relevant for SCZ; some of these small genomic regions mutations have already been modeled but much more work remains. 2D cultures derived from iPSCs can be used for differentiating into SCZ patient-specific neuronal subtypes but the lack of coherence between the different methods of iPSC reprogramming and differentiation can add confounding factors to the conclusions (Nayak et al. 2021). In addition, further methodological advancements in iPSC disease modeling are required to fully recapitulate the *in vivo* brain development system (Noh et al. 2017; Balan, Toyoshima, and Yoshikawa 2019).

SCZ management necessitates a better understanding of the pathophysiology and progression of the disease, as well as improved diagnostic methods for prevention and treatment (Chien et al. 2013). Transcriptomic studies in response to drug treatment in SCZ cohorts may enable finding associations with genomic signatures and biological markers that can eventually help in SCZ patient stratification and the development of precision medicine (Stern et al. 2018). Future iPSC studies should further utilize the knowledge that was gained by the genetic studies to classify SCZ patient samples into subtypes that are based on the clinical phenotypes to better understand the underlying pathophysiology and mechanisms that is specific to the SCZ disease spectrum.

## 4. METHODS

### 4.1 SCZ GWAS Genes analysis

To collect the comprehensive data of GWAS on SCZ subjects, we used GWAS Catalog (https://www.ebi.ac.uk/gwas/) (Buniello et al. 2018) and searched the database with the term “Schizophrenia” and downloaded the resulting association data table (.tsv file). This data was imported to Matlab for analysis. We used the column SNP_GENE_IDS which contains the Ensembl ID of the genes that contain SCZ-related rs (SNP-risk allele) within their sequence. Ensembl ID was matched with HGNC approved gene symbol using the method described below.

To count the number of publications for each ethnic group, we unified multiple studies from the same publication. The African sub-categories and the “Hispanic” category were unified into the “Hispanic or Latin American” category. For each gene, we determined the frequency by counting the number of times a gene variant was reported in a study with a p-value of <0.01. Note that one research article may include several studies (see the (Buniello et al. 2018) for details). This frequency was used to determine the font size in the word clouds in Figure 1 (b) and Figure 1 (d). Word-cloud figures were generated by Matlab’s word cloud function. Genes that are marked in red also appear in the list of 160 differentially expressed genes (DEGs) related to SCZ which was reported in the recent meta-analysis review (Merikangas et al. 2022).

### 4.2. Common genes between SCZ GWAS and SCZ DEGs

To intersect these two lists of genes, gene symbols were first standardized using HUGO Gene Nomenclature (https://www.genenames.org/) as described below and both the lists were searched for common genes among them and the word cloud was plotted using Matlab. The font size of the gene name is proportional to the number of times it has been reported as DEGs across different transcriptomic studies in SCZ clinical samples.

### 4.3. Standardization of Gene nomenclature

We downloaded the list of Ensembl ID and their matching symbol from biomart: https://www.ensembl.org/biomart/martview/. We chose the “Ensembl Genes 106“ database and the “Human Genes (GRCh38.p13)” dataset. We chose the attributes “Gene Stable ID” and “ Gene name” and downloaded the result CSV file. We also downloaded the database of HGNC Gene symbols from http://ftp.ebi.ac.uk/pub/databases/genenames/hgnc/tsv/hgnc_complete_set.txt which includes for each gene: approved gene symbol, previous gene symbols, alias gene symbols, and the Ensembl ID

For each Ensembl ID in the GWAS data, we did the following process to obtain its gene symbol. We searched for the ensemble ID both in the biomart results and in the HGNC database. If the Ensembl ID was found in the HGNC database we used the matching approved symbol from the HGNC database. If it didn’t match any ensemble ID in the HGNC database, we used the biomart symbol and compared it to the list of approved HGNC symbols. If the symbol was an approved HGNC symbol then we used it. If it was not an approved symbol, we searched it in the list of previous gene symbols in the HGNC database. If the symbol matched a previous symbol of a gene then we used the currently approved HGNC symbol of that gene (and not the previous symbol). This allowed matching HGNC approved nomenclature for all of the Ensembl IDs without ambiguity. However, three Ensembl IDs (ENSG00000145075, ENSG00000284548, ENSG00000284548) did not have a matching symbol either in biomart or in HGNC and were not used for the intersection. These three IDs are labeled as novel genes on the Ensembl website (using https://www.ensembl.org search). This process resulted in three gene symbols that were changed from their biomart symbol to the matching HGNC:

For the DEGs we did a similar process; if a gene symbol matched an approved HGNC gene symbol then we used it. If it did not match an approved HGNC gene, then we searched the previous gene names in the HGNC database and used the current symbol name. This process resulted in updating the following gene symbols:

### 4.4. Pie-chart distribution

All the pie charts were generated using Matlab from the collected data for respective figures. For Fig 1a and 1c, as mentioned above the respective data for ethnic groups in GWAS and the tissues used for transcriptomic studies were collected from the GWAS catalog and Merikangas et al.,2022. Fig1. a shows the number of publications that included each of the ethnic groups. Note that some papers included more than one ethnic group. For Fig 2a (i-iii), the pie charts show the distribution of data collected from approximately 52 papers, which used the iPSC-based model of SCZ. Note that for Fig 2a (iii), a single paper may include several cell types. The article list was generated from Pubmed using the search term “Schizophrenia iPSCs” from the timeline of 2011 to 2022. The relevant original research articles were referred and the data was collected for the “Type of tissue used” for reprogramming and “reprogramming methods” used for derivation of iPSCs as well as the “neural cell type” studied. We did not count approximately 8-10 articles for our analysis that had worked with previously derived iPSCs.

### 4.5. Heat maps

For Fig 1e, the differentially expressed miRNAs and lncRNAs from SCZ clinical samples including SCZ-iPSC-based models were listed. We analyzed for the presence of at least 2 reports for the same miRNAs and lncRNAs in any SCZ clinical sample. The list was converted to a heatmap using Matlab. The analysis did not include non-coding RNAs reported from animal models of SCZ. Also, not enough reports of circular RNAs in SCZ for metanalysis were available. Dark shade indicates the presence of differentially expressed non-coding RNAs in particular tissue and white indicates an absence of differentially expressed noncoding RNAs. Darker shades indicate more than one report. The color bar represents the frequency of differentially expressed non-coding RNAs across different publications.

For Fig 2b. The phenotypes reported from the SCZ-iPSC-related papers were listed and plotted into heat maps using Matlab. We also referred to data from the tables mentioned in (Räsänen et al. 2022). The blue color denotes the downregulation (↓) and the green upregulation (↑) of disease phenotypes. We have used arrows for simple representation. Both cortical glutamatergic neurons and induced glutamatergic neuron phenotypes have been unified in one cell type. A similar analysis was performed for GABAergic neurons. The color bar represents the frequency of phenotypes observed across different SCZ-iPSC-based models.

### 4.6. Tables

Table 1 and Table 2 are the list of genes for which the nomenclature was standardized according to HGNC approved symbols. Table 3 is based on previous literature reports on SCZ rare structural variants. We specifically also referred to the detailed meta-analysis report in (Rees et al. 2014) for rare structural variants reported for SCZ.

## APPENDIX 1

**Figure 1.**
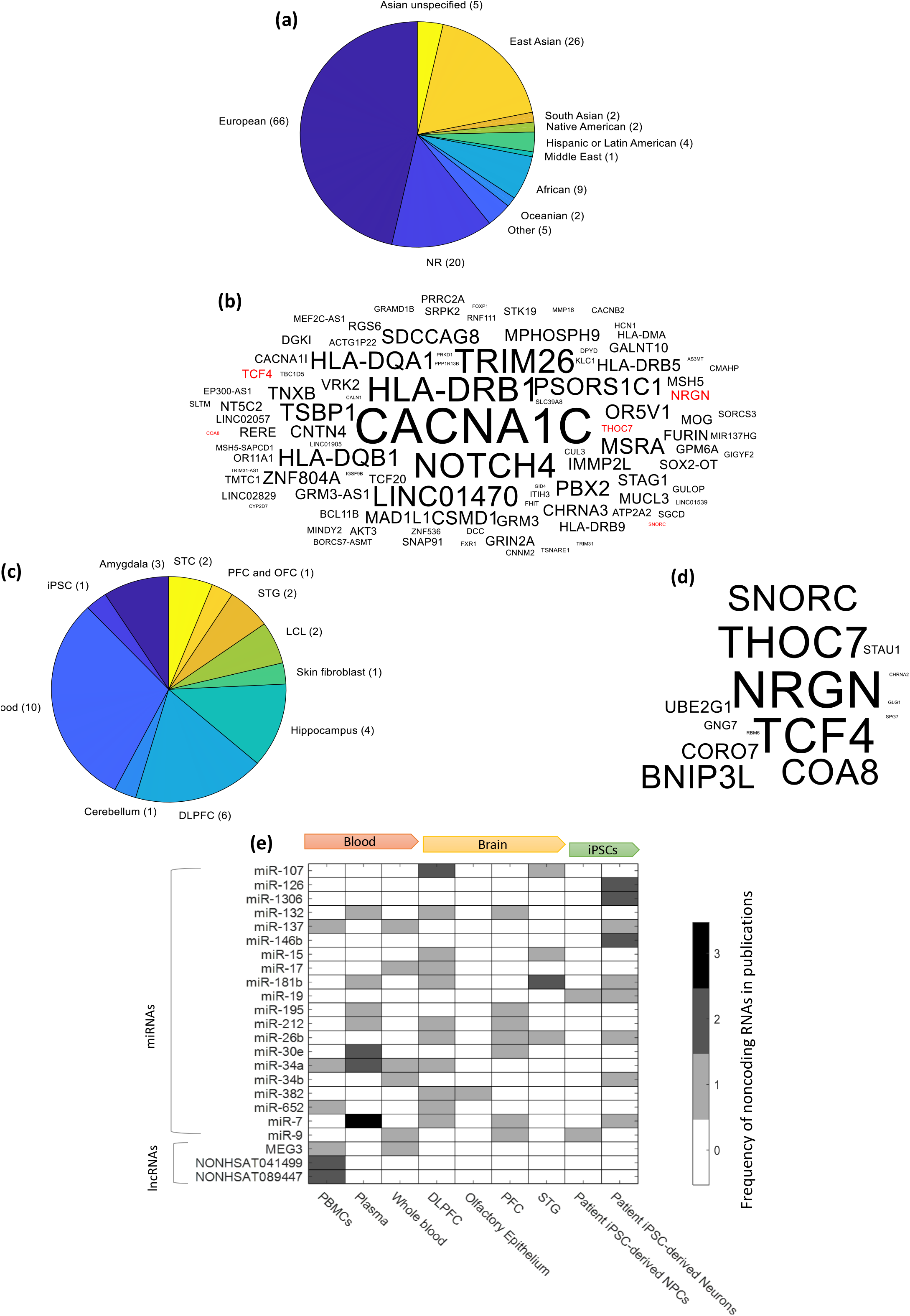
(a) A pie chart distribution of GWAS performed from different populations/ethnic groups across different GWAS publications. The plotted GWAS datasets were extracted from the GWAS catalog as mentioned in the methods. The pie chart includes only the discovery cohorts and no replication cohorts as described in the methods. (b) Genes significantly associated with SCZ according to the GWAS catalog. The font size is proportional to the frequency of the reported gene variant across different studies. Only studies which reported p-value < 0.01 for a specific variant were counted. Gene variants that were also reported as DEGs are colored in red. (c) A pie chart distribution of tissue types used in different transcriptomic studies on SCZ clinical samples. The list of studies was chosen based on the analysis done by Merikangas et al., 2022. For details, see the Methods section. LCL-lymphoblasts cell lines; PFC-Prefrontal cortex; DLPFC-Dorsolateral prefrontal cortex; STG-Superior temporal gyrus regions; OFC: orbitofrontal cortex. (d) A list of Common genes between Fig.1 (b) and highly reported DEGs across multiple transcriptomic studies on SCZ clinical samples (Merikangas et al., 2022). The font size of the gene is proportional to the frequency of the reported genes across different GWAS studies. (e) A heatmap showing shortlisted differentially expressed noncoding RNAs (miRNAs and lncRNAs) in different SCZ clinical tissues and patient-specific iPSCs. The list of miRNAs and lncRNAs was shortlisted based on the report of miRNAs and lncRNAs in at least 2 studies done on SCZ clinical samples including SCZ patient iPSCs. Dark shade indicates the presence of noncoding RNAs in particular tissue and white indicates the absence of noncoding RNAs. Darker shades indicate more than one report. PFC-Prefrontal cortex; DLPFC-Dorsolateral prefrontal cortex; STG-Superior temporal gyrus regions of postmortem brain. The color bar represents the frequency of non-coding RNAs across different publications. See the Methods section for more detail.

**Figure 2.**
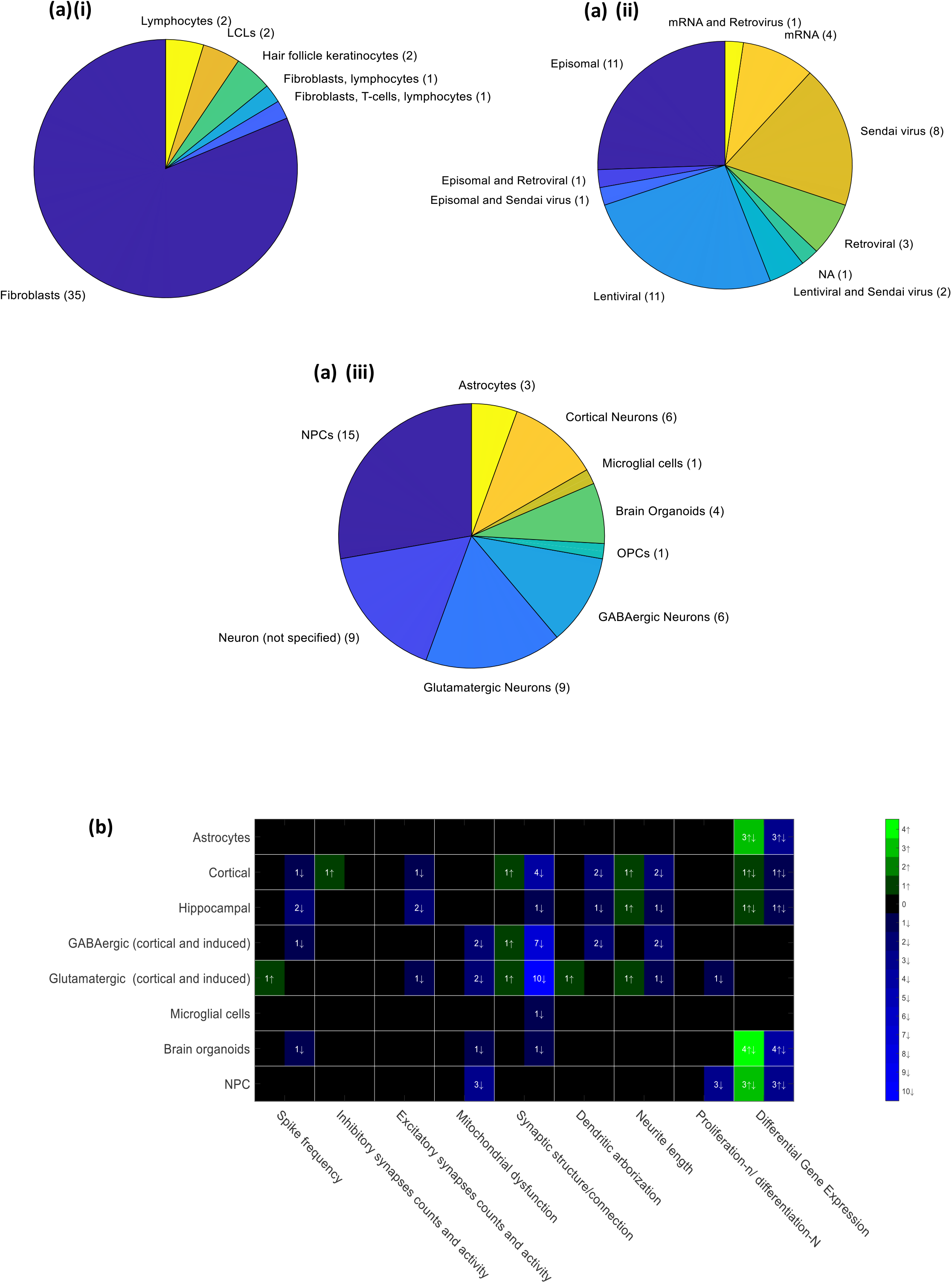
(a) A pie chart showing the distribution of i. Different clinical tissues are used for deriving iPSCs in different SCZ-iPSC-based publications. ii. The type of reprogramming method used for deriving SCZ patient-specific iPSCs in different publications. iii. Neural cell types studied in different SCZ-iPSC-based publications (b) A heatmap for the phenotypes observed during differentiation of SCZ-iPSCs into different cell types. The blue color denotes the downregulation (↓) and the green denotes an upregulation (↑) of disease phenotypes. No information or report is denoted by a black background. Differential gene expression observed in neural cell type has been denoted with both upregulation and downregulation (↑ ↓) symbol for DEGs and both blue and green colors has been used for DEGs. Both cortical glutamatergic neurons and induced glutamatergic neuron phenotypes have been unified in one cell type. A similar analysis has been done for GABAergic neurons. Proliferation-n stands for NPC proliferation while differentiation-N stands for neuronal subtype differentiation.

**Table 1.**
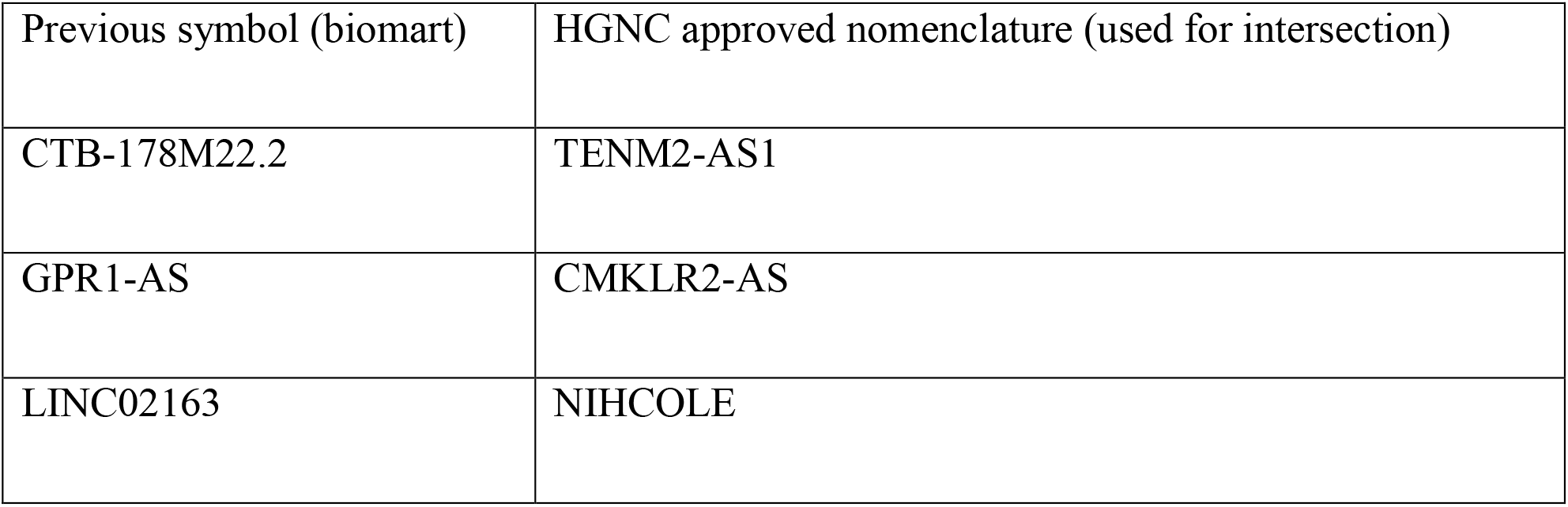
Gene variants from the GWAS database standardized according to HGNC approved nomenclature

| Previous symbol (biomart) | HGNC approved nomenclature (used for intersection) |
| --- | --- |
| CTB-178M22.2 | TENM2-AS1 |
| GPR1-AS | CMKLR2-AS |
| LINC02163 | NIHCOLE |

**Table 2.** DEGs are standardized according to the HGNC approved nomenclature

| Previous symbol | HGNC approved symbol (used for intersection) |
| --- | --- |
| C2orf82 | SNORC |
| MARCH2 | MARCHF2 |
| SMEK2 | PPP4R3B |
| APOPT1 | COA8 |
| C10orf54 | VSIR |
| C9orf16 | BBLN |
| CCDC130 | YJU2B |
| GPR56 | ADGRG1 |
| H1FX-AS1 | H1-10-AS1 |
| HIST1H2BD | H2BC5 |
| LINC00634 | SMIM45 |
| MARCH7 | MARCHF7 |
| SEPT5 | SEPTIN5 |

**Table 3.**
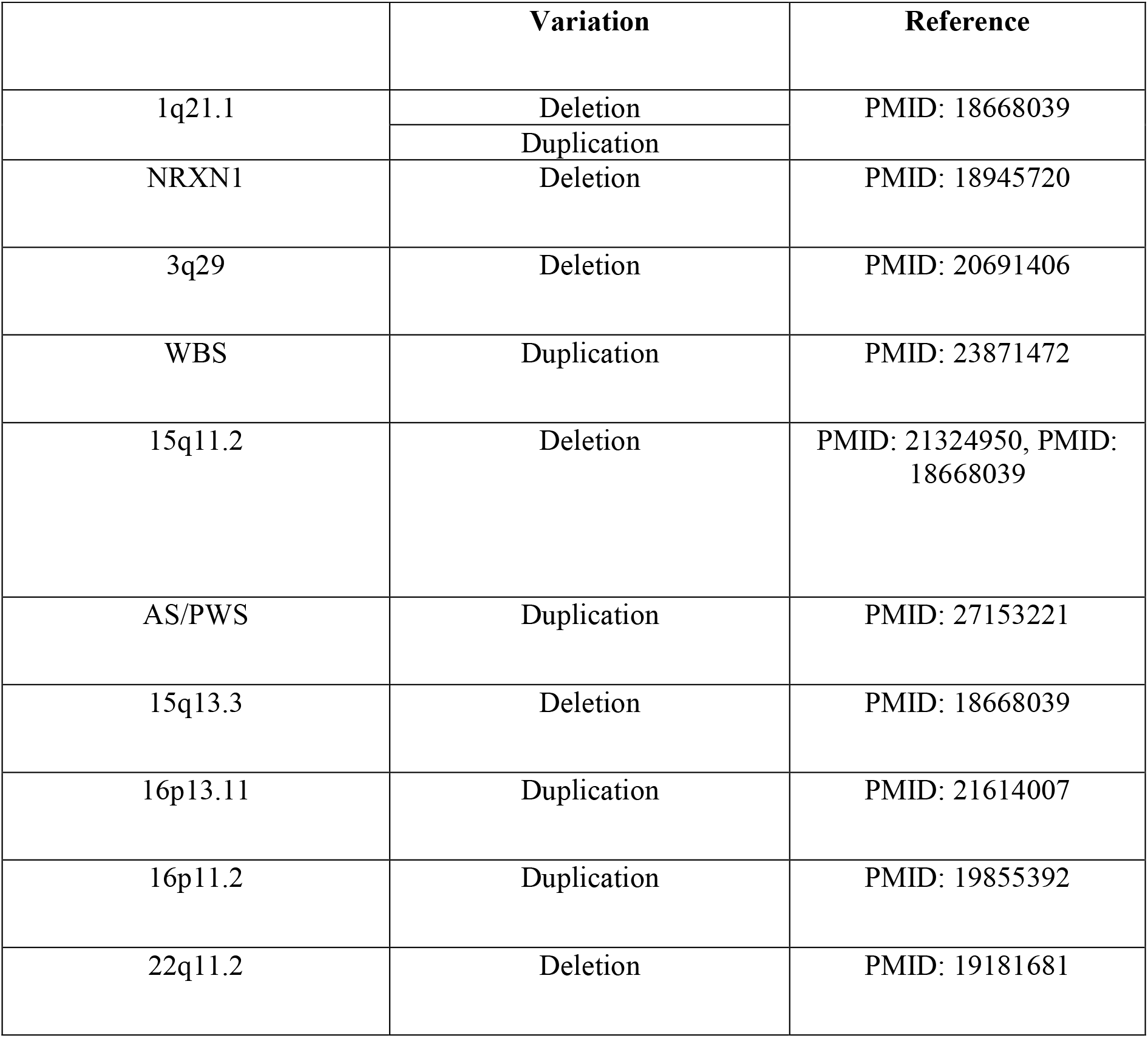
A list of eleven rare structural variants implicated in SCZ and reported to be significant in previous genetic meta-analysis studies (Rees et al. 2014). See methods for further details. ASD-Autism spectrum disease; ADHD-Attention deficit-hyperactivity disorder

## REFERENCES

Ahmad, Ruhel, Vincenza Sportelli, Michael Ziller, Dietmar Spengler, and Anke %J Cells Hoffmann. 2018. ’Tracing early neurodevelopment in schizophrenia with induced pluripotent stem cells’, 7: 140.

Akdeniz, Ceren, Heike Tost, Fabian Streit, Leila Haddad, Stefan Wüst Axel Schäfer, Michael Schneider, Marcella Rietschel, Peter Kirsch, and Andreas Meyer-Lindenberg. 2014. ‘Neuroimaging Evidence for a Role of Neural Social Stress Processing in Ethnic Minority– Associated Environmental Risk’, JAMA Psychiatry, 71: 672–80.

Akkouh, I. A., T. Hughes, V. M. Steen, J. C. Glover, O. A. Andreassen, S. Djurovic, and A. Szabo. 2021. ‘Transcriptome analysis reveals disparate expression of inflammation-related miRNAs and their gene targets in iPSC-astrocytes from people with schizophrenia’, Brain Behav Immun, 94: 235–44.

Akkouh, I. A., T. Ueland, L. Hansson, E. Inderhaug, T. Hughes, N. E. Steen, P. Aukrust, O. A. Andreassen, A. Szabo, and S. Djurovic. 2020. ‘Decreased IL-1β-induced CCL20 response in human iPSC-astrocytes in schizophrenia: Potential attenuating effects on recruitment of regulatory T cells’, Brain Behav Immun, 87: 634–44.

Almeida, Maria, Li Han, Marta Martin-Millan, Charles A. O’Brien, and Stavros C. Manolagas. 2007. ‘Oxidative Stress Antagonizes Wnt Signaling in Osteoblast Precursors by Diverting β-Catenin from T Cell Factor-to Forkhead Box O-mediated Transcription*’, Journal of Biological Chemistry, 282: 27298–305.

Balan, Shabeesh, Manabu Toyoshima, and Takeo %J Neurobiology of disease Yoshikawa. 2019. ’Contribution of induced pluripotent stem cell technologies to the understanding of cellular phenotypes in schizophrenia’, 131: 104162.

Barry, G., J. A. Briggs, D. P. Vanichkina, E. M. Poth, N. J. Beveridge, V. S. Ratnu, S. P. Nayler, K. Nones, J. Hu, T. W. Bredy, S. Nakagawa, F. Rigo, R. J. Taft, M. J. Cairns, S. Blackshaw, E. J. Wolvetang, and J. S. Mattick. 2014. ‘The long non-coding RNA Gomafu is acutely regulated in response to neuronal activation and involved in schizophrenia-associated alternative splicing’, Mol Psychiatry, 19: 486–94.

Bloom, Floyd E. 1993. ‘Advancing a Neurodevelopmental Origin for Schizophrenia’, Arch Gen Psychiatry, 50: 224–27.

Brant, Boris, Tchelet Stern, Huda Adwan Shekhidem, Liron Mizrahi, Idan Rosh, Yam Stern, Polina Ofer, Ayat Asleh, George K. Essien Umanah, Reem Jada, Nina S. Levy, Andrew P. Levy, and Shani Stern. 2021. ‘IQSEC2 mutation associated with epilepsy, intellectual disability, and autism results in hyperexcitability of patient-derived neurons and deficient synaptic transmission’, Mol Psychiatry, 26: 7498–508.

Brennand, K., J. N. Savas, Y. Kim, N. Tran, A. Simone, K. Hashimoto-Torii, K. G. Beaumont, H. J. Kim, A. Topol, I. Ladran, M. Abdelrahim, B. Matikainen-Ankney, S. h Chao, M. Mrksich, P. Rakic, G. Fang, B. Zhang, J. R. Yates, and F. H. Gage. 2015. ‘Phenotypic differences in hiPSC NPCs derived from patients with schizophrenia’, Mol Psychiatry, 20: 361–68.

Brennand, Kristen J., Anthony Simone, Jessica Jou, Chelsea Gelboin-Burkhart, Ngoc Tran, Sarah Sangar, Yan Li, Yangling Mu, Gong Chen, Diana Yu, Shane McCarthy, Jonathan Sebat, and Fred H. Gage. 2011. ‘Modelling schizophrenia using human induced pluripotent stem cells’, Nature, 473: 221–25.

Brown, Alan S, and Elena J %J American Journal of Psychiatry Derkits. 2010. ’Prenatal infection and schizophrenia: a review of epidemiologic and translational studies’, 167: 261–80.

Brown, Ellie, Richard Gray, Samantha Lo Monaco, Brian O’Donoghue, Barnaby Nelson, Andrew Thompson, Shona Francey, and Pat McGorry. 2020. ‘The potential impact of COVID-19 on psychosis: A rapid review of contemporary epidemic and pandemic research’, Schizophr Res, 222: 79–87.

Bundo, Miki, Manabu Toyoshima, Yohei Okada, Wado Akamatsu, Junko Ueda, Taeko Nemoto-Miyauchi, Fumiko Sunaga, Michihiro Toritsuka, Daisuke Ikawa, and Akiyoshi %J Neuron Kakita. 2014. ’Increased l1 retrotransposition in the neuronal genome in schizophrenia’, 81: 306–13.

Buniello, Annalisa, Jacqueline A L MacArthur, Maria Cerezo, Laura W Harris, James Hayhurst, Cinzia Malangone, Aoife McMahon, Joannella Morales, Edward Mountjoy, Elliot Sollis, Daniel Suveges, Olga Vrousgou, Patricia L Whetzel, Ridwan Amode, Jose A Guillen, Harpreet S Riat, Stephen J Trevanion, Peggy Hall, Heather Junkins, Paul Flicek, Tony Burdett, Lucia A Hindorff, Fiona Cunningham, and Helen Parkinson. 2018. ‘The NHGRI-EBI GWAS Catalog of published genome-wide association studies, targeted arrays and summary statistics 2019’, Nucleic Acids Research, 47: D1005–D12.

Callicott, Joseph H., Richard E. Straub, Lukas Pezawas, Michael F. Egan, Venkata S. Mattay, Ahmad R. Hariri, Beth A. Verchinski, Andreas Meyer-Lindenberg, Rishi Balkissoon, Bhaskar Kolachana, Terry E. Goldberg, and Daniel R. Weinberger. 2005. ’Variation in DISC1 affects hippocampal structure and function and increases risk for schizophrenia’, 102: 8627–32.

Cannon, M., P. B. Jones, and R. M. Murray. 2002. ‘Obstetric complications and schizophrenia: historical and meta-analytic review’, Am J Psychiatry, 159: 1080–92.

Cardno, A. G., and Gottesman, II. 2000. ‘Twin studies of schizophrenia: from bow-and-arrow concordances to star wars Mx and functional genomics’, Am J Med Genet, 97: 12–7.

Chacko, Mason, Asha Job, Fred Caston, Prem George, Adeeb Yacoub, and Ricardo Cáceda. 2020. ‘COVID-19-Induced Psychosis and Suicidal Behavior: Case Report’, SN Comprehensive Clinical Medicine, 2: 2391–95.

Chambers, Stuart M., Christopher A. Fasano, Eirini P. Papapetrou, Mark Tomishima, Michel Sadelain, and Lorenz Studer. 2009. ‘Highly efficient neural conversion of human ES and iPS cells by dual inhibition of SMAD signaling’, Nature biotechnology, 27: 275–80.

Chiang, C. H., Y. Su, Z. Wen, N. Yoritomo, C. A. Ross, R. L. Margolis, H. Song, and G. l Ming. 2011. ‘Integration-free induced pluripotent stem cells derived from schizophrenia patients with a DISC1 mutation’, Mol Psychiatry, 16: 358–60.

Chien, W. T., S. F. Leung, F. K. Yeung, and W. K. Wong. 2013. ‘Current approaches to treatments for schizophrenia spectrum disorders, part II: psychosocial interventions and patient-focused perspectives in psychiatric care’, Neuropsychiatr Dis Treat, 9: 1463–81.

Cleynen, Isabelle, Worrawat Engchuan, Matthew S. Hestand, Tracy Heung, Aaron M. Holleman, H. Richard Johnston, Thomas Monfeuga, Donna M. McDonald-McGinn, Raquel E. Gur, Bernice E. Morrow, Ann Swillen, Jacob A. S. Vorstman, Carrie E. Bearden, Eva W. C. Chow, Marianne van den Bree, Beverly S. Emanuel, Joris R. Vermeesch, Stephen T. Warren, Michael J. Owen, Pankaj Chopra, David J. Cutler, Richard Duncan, Alex V. Kotlar, Jennifer G. Mulle, Anna J. Voss, Michael E. Zwick, Alexander Diacou, Aaron Golden, Tingwei Guo, Jhih-Rong Lin, Tao Wang, Zhengdong Zhang, Yingjie Zhao, Christian Marshall, Daniele Merico, Andrea Jin, Brenna Lilley, Harold I. Salmons, Oanh Tran, Peter Holmans, Antonio Pardinas, James T. R. Walters, Wolfram Demaerel, Erik Boot, Nancy J. Butcher, Gregory A. Costain, Chelsea Lowther, Rens Evers, Therese A. M. J. van Amelsvoort, Esther van Duin, Claudia Vingerhoets, Jeroen Breckpot, Koen Devriendt, Elfi Vergaelen, Annick Vogels, T. Blaine Crowley, Daniel E. McGinn, Edward M. Moss, Robert J. Sharkus, Marta Unolt, Elaine H. Zackai, Monica E. Calkins, Robert S. Gallagher, Ruben C. Gur, Sunny X. Tang, Rosemarie Fritsch, Claudia Ornstein, Gabriela M. Repetto, Elemi Breetvelt, Sasja N. Duijff, Ania Fiksinski, Hayley Moss, Maria Niarchou, Kieran C. Murphy, Sarah E. Prasad, Eileen M. Daly, Maria Gudbrandsen, Clodagh M. Murphy, Declan G. Murphy, Antonio Buzzanca, Fabio Di Fabio, Maria C. Digilio, Maria Pontillo, Bruno Marino, Stefano Vicari, Karlene Coleman, Joseph F. Cubells, Opal Y. Ousley, Miri Carmel, Doron Gothelf, Ehud Mekori-Domachevsky, Elena Michaelovsky, Ronnie Weinberger, Abraham Weizman, Leila Kushan, Maria Jalbrzikowski, Marco Armando, Stéphan Eliez, Corrado Sandini, Maude Schneider, Frédérique Sloan Béna, Kevin M. Antshel, Wanda Fremont, Wendy R. Kates, Raoul Belzeaux, Tiffany Busa, Nicole Philip, Linda E. Campbell, Kathryn L. McCabe, Stephen R. Hooper, Kelly Schoch, Vandana Shashi, Tony J. Simon, Flora Tassone, Celso Arango, David Fraguas, Sixto García-Miñaúr, Jaume Morey-Canyelles, Jordi Rosell, Damià H. Suñer, Jasna Raventos-Simic, Michael P. Epstein, Nigel M. Williams, Anne S. Bassett, D. S. Brain International 22q, and Consortium Behavior. 2021. ‘Genetic contributors to risk of schizophrenia in the presence of a 22q11.2 deletion’, Mol Psychiatry, 26: 4496–510.

Costain, G., A. C. Lionel, D. Merico, P. Forsythe, K. Russell, C. Lowther, T. Yuen, J. Husted, D. J. Stavropoulos, M. Speevak, E. W. Chow, C. R. Marshall, S. W. Scherer, and A. S. Bassett. 2013. ‘Pathogenic rare copy number variants in community-based schizophrenia suggest a potential role for clinical microarrays’, Hum Mol Genet, 22: 4485–501.

D’Aiuto, Leonardo, Konasale M. Prasad, Catherine H. Upton, Luigi Viggiano, Jadranka Milosevic, Giorgio Raimondi, Lora McClain, Kodavali Chowdari, Jay Tischfield, Michael Sheldon, Jennifer C. Moore, Robert H. Yolken, Paul R. Kinchington, and Vishwajit L. Nimgaonkar. 2014. ‘Persistent Infection by HSV-1 Is Associated With Changes in Functional Architecture of iPSC-Derived Neurons and Brain Activation Patterns Underlying Working Memory Performance’, Schizophr Bull, 41: 123–32.

Da Silveira Paulsen Bruna, Renata De Moraes Maciel, Antonio Galina, Mariana Souza Da Silveira, Cleide Dos Santos Souza, Hannah Drummond, Ernesto Nascimento Pozzatto, Hamilton Silva Junior, Leonardo Chicaybam, Raffael Massuda, Pedro Setti-Perdigão, Martin Bonamino, Paulo Silva Belmonte-De-Abreu, Newton Gonçalves Castro, Helena Brentani, and Stevens Kastrup Rehen. 2012. ’Altered Oxygen Metabolism Associated to Neurogenesis of Induced Pluripotent Stem Cells Derived from a Schizophrenic Patient’, 21: 1547–59.

De Hert, M., J. Detraux, R. van Winkel, W. Yu, and C. U. Correll. 2011. ‘Metabolic and cardiovascular adverse effects associated with antipsychotic drugs’, Nat Rev Endocrinol, 8: 114–26.

De Los Angeles, Alejandro, Michael B. Fernando, Nicola A. L. Hall, Kristen J. Brennand, Paul J. Harrison, Brady J. Maher, Daniel R. Weinberger, and Elizabeth M. Tunbridge. 2021. ‘Induced Pluripotent Stem Cells in Psychiatry: An Overview and Critical Perspective’, Biological Psychiatry, 90: 362–72.

Doyle, Glenn A., Richard C. Crist, Emre T. Karatas, Matthew J. Hammond, Adam D. Ewing, Thomas N. Ferraro, Chang-Gyu Hahn, and Wade H. Berrettini. 2017. ‘Analysis of LINE-1 Elements in DNA from Postmortem Brains of Individuals with Schizophrenia’, Neuropsychopharmacology : official publication of the American College of Neuropsychopharmacology, 42: 2602–11.

Dror, N., E. Klein, R. Karry, A. Sheinkman, Z. Kirsh, M. Mazor, M. Tzukerman, and D. Ben-Shachar. 2002. ‘State-dependent alterations in mitochondrial complex I activity in platelets: a potential peripheral marker for schizophrenia’, Mol Psychiatry, 7: 995–1001.

Estes, Myka L, and A Kimberley %J Science McAllister. 2016. ’Maternal immune activation: Implications for neuropsychiatric disorders’, 353: 772–77.

Farrell, M. S., T. Werge, P. Sklar, M. J. Owen, R. A. Ophoff, M. C. O’Donovan, A. Corvin, S. Cichon, and P. F. Sullivan. 2015. ‘Evaluating historical candidate genes for schizophrenia’, Mol Psychiatry, 20: 555–62.

Fatemi, S. Hossein, and Timothy D. Folsom. 2009. ‘The Neurodevelopmental Hypothesis of Schizophrenia, Revisited’, Schizophr Bull, 35: 528–48.

Haijma, S. V., N. Van Haren, W. Cahn, P. C. Koolschijn, H. E. Hulshoff Pol, and R. S. Kahn. 2013. ‘Brain volumes in schizophrenia: a meta-analysis in over 18 000 subjects’, Schizophr Bull, 39: 1129–38.

Harb, Hani, Mehdi Benamar, Peggy S. Lai, Paola Contini, Jason W. Griffith, Elena Crestani, Klaus Schmitz-Abe, Qian Chen, Jason Fong, Luca Marri, Gilberto Filaci, Genny Del Zotto, Novalia Pishesha, Stephen Kolifrath, Achille Broggi, Sreya Ghosh, Metin Yusuf Gelmez, Fatma Betul Oktelik, Esin Aktas Cetin, Ayca Kiykim, Murat Kose, Ziwei Wang, Ye Cui, Xu G. Yu, Jonathan Z. Li, Lorenzo Berra, Emmanuel Stephen-Victor, Louis-Marie Charbonnier, Ivan Zanoni, Hidde Ploegh, Gunnur Deniz, Raffaele De Palma, and Talal A. Chatila. 2021. ‘Notch4 signaling limits regulatory T-cell-mediated tissue repair and promotes severe lung inflammation in viral infections’, Immunity, 54: 1186–99.e7.

Hathy, Edit, Eszter Szabó, Nóra Varga, Zsuzsa Erdei, Csongor Tordai, Boróka Czehlár, Máté Baradits, Bálint Jezsó, Júlia Koller, László %J Stem cell research Nagy, and therapy. 2020. ’Investigation of de novo mutations in a schizophrenia case-parent trio by induced pluripotent stem cell-based in vitro disease modeling: convergence of schizophrenia-and autism-related cellular phenotypes’, 11: 1–15.

Henriksen, Mads G., Julie Nordgaard, and Lennart B. Jansson. 2017. ’Genetics of Schizophrenia: Overview of Methods, Findings and Limitations’, 11.

Indelicato, Elisabetta, and Sylvia Boesch. 2021. ’From Genotype to Phenotype: Expanding the Clinical Spectrum of CACNA1A Variants in the Era of Next Generation Sequencing’, 12.

Irish Schizophrenia Genomics, Consortium, and Consortium the Wellcome Trust Case Control. 2012. ‘Genome-wide association study implicates HLA-C*01:02 as a risk factor at the major histocompatibility complex locus in schizophrenia’, Biological Psychiatry, 72: 620–28.

Ishii, T., M. Ishikawa, K. Fujimori, T. Maeda, I. Kushima, Y. Arioka, D. Mori, Y. Nakatake, B. Yamagata, S. Nio, T. A. Kato, N. Yang, M. Wernig, S. Kanba, M. Mimura, N. Ozaki, and H. Okano. 2019. ‘In Vitro Modeling of the Bipolar Disorder and Schizophrenia Using Patient-Derived Induced Pluripotent Stem Cells with Copy Number Variations of PCDH15 and RELN’, eNeuro, 6.

Iwamoto, Kazuya, Miki Bundo, and Tadafumi Kato. 2004. ‘Altered expression of mitochondria-related genes in postmortem brains of patients with bipolar disorder or schizophrenia, as revealed by large-scale DNA microarray analysis’, Human Molecular Genetics, 14: 241–53.

Iyegbe, Conrad, Desmond Campbell, Amy Butler, Olesya Ajnakina, and Pak Sham. 2014. ‘The emerging molecular architecture of schizophrenia, polygenic risk scores and the clinical implications for GxE research’, Social Psychiatry and Psychiatric Epidemiology, 49: 169–82.

Kahn, René S., Iris E. Sommer, Robin M. Murray, Andreas Meyer-Lindenberg, Daniel R. Weinberger, Tyrone D. Cannon, Michael O’Donovan, Christoph U. Correll, John M. Kane, Jim van Os, and Thomas R. Insel. 2015. ‘Schizophrenia’, Nature Reviews Disease Primers, 1: 15067.

Kaiser, T., Y. Zhou, and G. Feng. 2017. ‘Animal models for neuropsychiatric disorders: prospects for circuit intervention’, Curr Opin Neurobiol, 45: 59–65.

Kamat, Pradip K., Anuradha Kalani, Shivika Rai, Supriya Swarnkar, Santoshkumar Tota, Chandishwar Nath, and Neetu Tyagi. 2016. ‘Mechanism of Oxidative Stress and Synapse Dysfunction in the Pathogenesis of Alzheimer’s Disease: Understanding the Therapeutics Strategies’, Molecular neurobiology, 53: 648–61.

Kane, J. M., and C. U. Correll. 2010. ‘Past and present progress in the pharmacologic treatment of schizophrenia’, J Clin Psychiatry, 71: 1115–24.

Kathuria, Annie, Kara Lopez-Lengowski, Smita S Jagtap, Donna McPhie, Roy H Perlis, Bruce M Cohen, and Rakesh %J JAMA psychiatry Karmacharya. 2020. ’Transcriptomic landscape and functional characterization of induced pluripotent stem cell–derived cerebral organoids in schizophrenia’, 77: 745–54.

Kathuria, Annie, Kara Lopez-Lengowski, Bradley Watmuff, Donna McPhie, Bruce M Cohen, and Rakesh %J Translational psychiatry Karmacharya. 2019a. ’Synaptic deficits in iPSC-derived cortical interneurons in schizophrenia are mediated by NLGN2 and rescued by N-acetylcysteine’, 9: 1–13.

Kathuria, Annie, Kara Lopez-Lengowski, Bradley Watmuff, Donna McPhie, Bruce M. Cohen, and Rakesh Karmacharya. 2019b. ‘Synaptic deficits in iPSC-derived cortical interneurons in schizophrenia are mediated by NLGN2 and rescued by N-acetylcysteine’, Transl Psychiatry, 9: 321.

Kohn, R., S. Saxena, I. Levav, and B. Saraceno. 2004. ‘The treatment gap in mental health care’, Bull World Health Organ, 82: 858–66.

Lam, Max, Chia-Yen Chen, Zhiqiang Li, Alicia R. Martin, Julien Bryois, Xixian Ma, Helena Gaspar, Masashi Ikeda, Beben Benyamin, Brielin C. Brown, Ruize Liu, Wei Zhou, Lili Guan, Yoichiro Kamatani, Sung-Wan Kim, Michiaki Kubo, Agung A. A. A. Kusumawardhani, Chih-Min Liu, Hong Ma, Sathish Periyasamy, Atsushi Takahashi, Zhida Xu, Hao Yu, Feng Zhu, Wei J. Chen, Stephen Faraone, Stephen J. Glatt, Lin He, Steven E. Hyman, Hai-Gwo Hwu, Steven A. McCarroll, Benjamin M. Neale, Pamela Sklar, Dieter B. Wildenauer, Xin Yu, Dai Zhang, Bryan J. Mowry, Jimmy Lee, Peter Holmans, Shuhua Xu, Patrick F. Sullivan, Stephan Ripke, Michael C. O’Donovan, Mark J. Daly, Shengying Qin, Pak Sham, Nakao Iwata, Kyung S. Hong, Sibylle G. Schwab, Weihua Yue, Ming Tsuang, Jianjun Liu, Xiancang Ma, René S. Kahn, Yongyong Shi, Hailiang Huang, Consortium Schizophrenia Working Group of the Psychiatric Genomics, Consortium Indonesia Schizophrenia, REsearch on schizophreniA neTwork-China Genetic, and Netherlands the. 2019. ‘Comparative genetic architectures of schizophrenia in East Asian and European populations’, Nature Genetics, 51: 1670–78.

Lederbogen, Florian, Peter Kirsch, Leila Haddad, Fabian Streit, Heike Tost, Philipp Schuch, Stefan Wüst, Jens C. Pruessner, Marcella Rietschel, Michael Deuschle, and Andreas Meyer-Lindenberg. 2011. ‘City living and urban upbringing affect neural social stress processing in humans’, Nature, 474: 498–501.

Lee, E. E., J. Liu, X. Tu, B. W. Palmer, L. T. Eyler, and D. V. Jeste. 2018. ‘A widening longevity gap between people with schizophrenia and general population: A literature review and call for action’, Schizophr Res, 196: 9–13.

Leon, J., K. Sakumi, E. Castillo, Z. Sheng, S. Oka, and Y. Nakabeppu. 2016. ‘8-Oxoguanine accumulation in mitochondrial DNA causes mitochondrial dysfunction and impairs neuritogenesis in cultured adult mouse cortical neurons under oxidative conditions’, Sci Rep, 6: 22086.

Leucht, S., A. Cipriani, L. Spineli, D. Mavridis, D. Orey, F. Richter, M. Samara, C. Barbui, R. R. Engel, J. R. Geddes, W. Kissling, M. P. Stapf, B. Lässig, G. Salanti, and J. M. Davis. 2013. ‘Comparative efficacy and tolerability of 15 antipsychotic drugs in schizophrenia: a multiple-treatments meta-analysis’, Lancet, 382: 951–62.

Li, Jianping, Sean K. Ryan, Erik Deboer, Kieona Cook, Shane Fitzgerald, Herbert M. Lachman, Douglas C. Wallace, Ethan M. Goldberg, and Stewart A. Anderson. 2019. ‘Mitochondrial deficits in human iPSC-derived neurons from patients with 22q11.2 deletion syndrome and schizophrenia’, Transl Psychiatry, 9: 302.

Liu, Y., X. Chang, C. G. Hahn, R. E. Gur, P. A. M. Sleiman, and H. Hakonarson. 2018. ‘Non-coding RNA dysregulation in the amygdala region of schizophrenia patients contributes to the pathogenesis of the disease’, Transl Psychiatry, 8: 44.

Maher, B. 2008. ‘Personal genomes: The case of the missing heritability’, Nature, 456: 18–21.

Mahmoudi, Ebrahim, Chantel Fitzsimmons, Michael P. Geaghan, Cynthia Shannon Weickert, Joshua R. Atkins, Xi Wang, and Murray J. Cairns. 2019. ‘Circular RNA biogenesis is decreased in postmortem cortical gray matter in schizophrenia and may alter the bioavailability of associated miRNA’, Neuropsychopharmacology : official publication of the American College of Neuropsychopharmacology, 44: 1043–54.

Malaspina, D., S. Harlap, S. Fennig, D. Heiman, D. Nahon, D. Feldman, and E. S. Susser. 2001. ‘Advancing paternal age and the risk of schizophrenia’, Arch Gen Psychiatry, 58: 361–7.

Massaad, Cynthia A., and Eric Klann. 2011. ‘Reactive oxygen species in the regulation of synaptic plasticity and memory’, Antioxidants & redox signaling, 14: 2013–54.

Meltzer, H. Y., L. Alphs, A. I. Green, A. C. Altamura, R. Anand, A. Bertoldi, M. Bourgeois, G. Chouinard, M. Z. Islam, J. Kane, R. Krishnan, J. P. Lindenmayer, and S. Potkin. 2003. ‘Clozapine treatment for suicidality in schizophrenia: International Suicide Prevention Trial (InterSePT)’, Arch Gen Psychiatry, 60: 82–91.

Merikangas, Alison K., Matthew Shelly, Alexys Knighton, Nicholas Kotler, Nicole Tanenbaum, and Laura Almasy. 2022. ‘What genes are differentially expressed in individuals with schizophrenia? A systematic review’, Mol Psychiatry, 27: 1373–83.

Mutch, Christopher A, Nobuo Funatsu, Edwin S Monuki, and Anjen %J Journal of Neuroscience Chenn. 2009. ’β-catenin signaling levels in progenitors influence the laminar cell fates of projection neurons’, 29: 13710–19.

Narla, ST, YW Lee, CA Benson, P Sarder, KJ Brennand, EK Stachowiak, and MK %J Schizophrenia research Stachowiak. 2017. ’Common developmental genome deprogramming in schizophrenia—Role of Integrative Nuclear FGFR1 Signaling (INFS)’, 185: 17–32.

Nayak, Ritu, Idan Rosh, Irina Kustanovich, and Shani Stern. 2021. ’Mood Stabilizers in Psychiatric Disorders and Mechanisms Learnt from In Vitro Model Systems’, 22: 9315.

Nestler, E. J., and S. E. Hyman. 2010. ‘Animal models of neuropsychiatric disorders’, Nat Neurosci, 13: 1161–9.

Ni, Peiyan, Haneul Noh, Gun-Hoo Park, Zhicheng Shao, Youxin Guan, James M Park, Sophy Yu, Joy S Park, Joseph T Coyle, and Daniel R %J Molecular psychiatry Weinberger. 2020. ’iPSC-derived homogeneous populations of developing schizophrenia cortical interneurons have compromised mitochondrial function’, 25: 2873–88.

Nielsen, R. E., S. Levander, G. Kjaersdam Telléus, S. O. Jensen, T. Østergaard Christensen, and S. Leucht. 2015. ‘Second-generation antipsychotic effect on cognition in patients with schizophrenia--a meta-analysis of randomized clinical trials’, Acta Psychiatr Scand, 131: 185–96.

Noh, H., Z. Shao, J. T. Coyle, and S. Chung. 2017. ‘Modeling schizophrenia pathogenesis using patient-derived induced pluripotent stem cells (iPSCs)’, Biochim Biophys Acta Mol Basis Dis, 1863: 2382–87.

Notaras, Michael, Aiman Lodhi, Haoyun Fang, David Greening, and Dilek %J Translational Psychiatry Colak. 2021. ’The proteomic architecture of schizophrenia iPSC-derived cerebral organoids reveals alterations in GWAS and neuronal development factors’, 11: 1–16.

Page, Stephanie Cerceo, Srinidhi Rao Sripathy, Federica Farinelli, Zengyou Ye, Yanhong Wang, Daniel J. Hiler, Elizabeth A. Pattie, Claudia V. Nguyen, Madhavi Tippani, Rebecca L. Moses, Huei-Ying Chen, Matthew Nguyen Tran, Nicholas J. Eagles, Joshua M. Stolz, Joseph L. Catallini, Olivia R. Soudry, Dwight Dickinson, Karen F. Berman, Jose A. Apud, Daniel R. Weinberger, Keri Martinowich, Andrew E. Jaffe, Richard E. Straub, and Brady J. Maher. 2022. ’Electrophysiological measures from human iPSC-derived neurons are associated with schizophrenia clinical status and predict individual cognitive performance’, 119: e2109395119.

Pak, ChangHui, Tamas Danko, Vincent R. Mirabella, Jinzhao Wang, Yingfei Liu, Madhuri Vangipuram, Sarah Grieder, Xianglong Zhang, Thomas Ward, Yu-Wen Alvin Huang, Kang Jin, Philip Dexheimer, Eric Bardes, Alexis Mitelpunkt, Junyi Ma, Michael McLachlan, Jennifer C. Moore, Pingping Qu, Carolin Purmann, Jeffrey L. Dage, Bradley J. Swanson, Alexander E. Urban, Bruce J. Aronow, Zhiping P. Pang, Douglas F. Levinson, Marius Wernig, and Thomas C. Südhof. 2021. ’Cross-platform validation of neurotransmitter release impairments in schizophrenia patient-derived <i>NRXN1</i>-mutant neurons’, 118: e2025598118.

Panaccione, I., F. Napoletano, A. M. Forte, G. D. Kotzalidis, A. Del Casale, C. Rapinesi, C. Brugnoli, D. Serata, F. Caccia, I. Cuomo, E. Ambrosi, A. Simonetti, V. Savoja, L. De Chiara, E. Danese, G. Manfredi, D. Janiri, M. Motolese, F. Nicoletti, P. Girardi, and G. Sani. 2013. ‘Neurodevelopment in schizophrenia: the role of the wnt pathways’, Curr Neuropharmacol, 11: 535–58.

Park, Gun-Hoo, Haneul Noh, Zhicheng Shao, Peiyan Ni, Yiren Qin, Dongxin Liu, Cameron P Beaudreault, Joy S Park, Chiderah P Abani, and James M %J Nature neuroscience Park. 2020. ’Activated microglia cause metabolic disruptions in developmental cortical interneurons that persist in interneurons from individuals with schizophrenia’, 23: 1352–64.

Pedrosa, Erika, Vladislav Sandler, Abhishek Shah, Reed Carroll, Chanjung Chang, Shira Rockowitz, Xingyi Guo, Deyou Zheng, and Herbert M. Lachman. 2011. ‘Development of Patient-Specific Neurons in Schizophrenia Using Induced Pluripotent Stem Cells’, Journal of Neurogenetics, 25: 88–103.

Pinnapureddy, Ashish R., Cherie Stayner, John McEwan, Olivia Baddeley, John Forman, and Michael R. Eccles. 2015. ‘Large animal models of rare genetic disorders: sheep as phenotypically relevant models of human genetic disease’, Orphanet Journal of Rare Diseases, 10: 107.

Potkin, Steven G., John M. Kane, Christoph U. Correll, Jean-Pierre Lindenmayer, Ofer Agid, Stephen R. Marder, Mark Olfson, and Oliver D. Howes. 2020. ‘The neurobiology of treatment-resistant schizophrenia: paths to antipsychotic resistance and a roadmap for future research’, npj Schizophrenia, 6: 1.

Prabakaran, S., J. E. Swatton, M. M. Ryan, S. J. Huffaker, Jt-J. Huang, J. L. Griffin, M. Wayland, T. Freeman, F. Dudbridge, K. S. Lilley, N. A. Karp, S. Hester, D. Tkachev, M. L. Mimmack, R. H. Yolken, M. J. Webster, E. F. Torrey, and S. Bahn. 2004. ‘Mitochondrial dysfunction in schizophrenia: evidence for compromised brain metabolism and oxidative stress’, Mol Psychiatry, 9: 684–97.

Purcell, Shaun M., Naomi R. Wray, Jennifer L. Stone, Peter M. Visscher, Michael C. O’Donovan, Patrick F. Sullivan, Pamela Sklar, Shaun M. Purcell, Jennifer L. Stone, Patrick F. Sullivan, Douglas M. Ruderfer, Andrew McQuillin, Derek W. Morris, Colm T. O’Dushlaine, Aiden Corvin, Peter A. Holmans, Michael C. O’Donovan, Pamela Sklar, Naomi R. Wray, Stuart Macgregor, Pamela Sklar, Patrick F. Sullivan, Michael C. O’Donovan, Peter M. Visscher, Hugh Gurling, Douglas H. R. Blackwood, Aiden Corvin, Nick J. Craddock, Michael Gill, Christina M. Hultman, George K. Kirov, Paul Lichtenstein, Andrew McQuillin, Walter J. Muir, Michael C. O’Donovan, Michael J. Owen, Carlos N. Pato, Shaun M. Purcell, Edward M. Scolnick, David St Clair, Jennifer L. Stone, Patrick F. Sullivan, Pamela Sklar, Michael C. O’Donovan, George K. Kirov, Nick J. Craddock, Peter A. Holmans, Nigel M. Williams, Lyudmila Georgieva, Ivan Nikolov, N. Norton, H. Williams, Draga Toncheva, Vihra Milanova, Michael J. Owen, Christina M. Hultman, Paul Lichtenstein, Emma F. Thelander, Patrick Sullivan, Derek W. Morris, Colm T. O’Dushlaine, Elaine Kenny, Emma M. Quinn, Michael Gill, Aiden Corvin, Andrew McQuillin, Khalid Choudhury, Susmita Datta, Jonathan Pimm, Srinivasa Thirumalai, Vinay Puri, Robert Krasucki, Jacob Lawrence, Digby Quested, Nicholas Bass, Hugh Gurling, Caroline Crombie, Gillian Fraser, Soh Leh Kuan, Nicholas Walker, David St Clair, Douglas H. R. Blackwood, Walter J. Muir, Kevin A. McGhee, Ben Pickard, Pat Malloy, Alan W. Maclean, Margaret Van Beck, Naomi R. Wray, Stuart Macgregor, Peter M. Visscher, Michele T. Pato, Helena Medeiros, Frank Middleton, Celia Carvalho, Christopher Morley, Ayman Fanous, David Conti, James A. Knowles, Carlos Paz Ferreira, Antonio Macedo, M. Helena Azevedo, Carlos N. Pato, Jennifer L. Stone, Douglas M. Ruderfer, Andrew N. Kirby, Manuel A. R. Ferreira, Mark J. Daly, Shaun M. Purcell, Pamela Sklar, Shaun M. Purcell, Jennifer L. Stone, Kimberly Chambert, Douglas M. Ruderfer, Finny Kuruvilla, Stacey B. Gabriel, Kristin Ardlie, Jennifer L. Moran, Mark J. Daly, Edward M. Scolnick, Pamela Sklar, Consortium The International Schizophrenia, preparation Manuscript, analysis Data, Gwas analysis subgroup, subgroup Polygene analyses, committee Management, University Cardiff, Hill Karolinska Institutet/University of North Carolina at Chapel, Dublin Trinity College, London University College, Aberdeen University of, Edinburgh University of, Research Queensland Institute of Medical, California University of Southern, Hospital Massachusetts General, Research Stanley Center for Psychiatric, M. I. T. Broad Institute of, and Harvard. 2009. ‘Common polygenic variation contributes to risk of schizophrenia and bipolar disorder’, Nature, 460: 748–52.

Quraishi, Imran H., Shani Stern, Kile P. Mangan, Yalan Zhang, Syed R. Ali, Michael R. Mercier, Maria C. Marchetto, Michael J. McLachlan, Eugenia M. Jones, Fred H. Gage, and Leonard K. Kaczmarek. 2019. ’An Epilepsy-Associated KCNT1 Mutation Enhances Excitability of Human iPSC-Derived Neurons by Increasing Slack K<sub>Na</sub> Currents’, 39: 7438–49.

Qureshi, Irfan A., and Mark F. Mehler. 2012. ‘Emerging roles of non-coding RNAs in brain evolution, development, plasticity and disease’, Nature Reviews Neuroscience, 13: 528–41.

Räsänen, Noora, Jari Tiihonen, Marja Koskuvi, Šárka Lehtonen, and Jari %J Trends in neurosciences Koistinaho. 2022. ’The iPSC perspective on schizophrenia’, 45: 8–26.

Rees, Elliott, James T. R. Walters, Lyudmila Georgieva, Anthony R. Isles, Kimberly D. Chambert, Alexander L. Richards, Gerwyn Mahoney-Davies, Sophie E. Legge, Jennifer L. Moran, Steven A. McCarroll, Michael C. O’Donovan, Michael J. Owen, and George Kirov. 2014. ‘Analysis of copy number variations at 15 schizophrenia-associated loci’, The British journal of psychiatry : the journal of mental science, 204: 108–14.

Rhoades, Raina, Fatimah Jackson, and Shaolei Teng. 2019. ‘Discovery of rare variants implicated in schizophrenia using next-generation sequencing’, Journal of Translational Genetics and Genomics, 3: 1.

Ripke, S., C. O’Dushlaine, K. Chambert, J. L. Moran, A. K. Kahler, S. Akterin, S. E. Bergen, A. L. Collins, J. J. Crowley, M. Fromer, Y. Kim, S. H. Lee, P. K. Magnusson, N. Sanchez, E. A. Stahl, S. Williams, N. R. Wray, K. Xia, F. Bettella, A. D. Borglum, B. K. Bulik-Sullivan, P. Cormican, N. Craddock, C. de Leeuw, N. Durmishi, M. Gill, V. Golimbet, M. L. Hamshere, P. Holmans, D. M. Hougaard, K. S. Kendler, K. Lin, D. W. Morris, O. Mors, P. B. Mortensen, B. M. Neale, F. A. O’Neill, M. J. Owen, M. P. Milovancevic, D. Posthuma, J. Powell, A. L. Richards, B. P. Riley, D. Ruderfer, D. Rujescu, E. Sigurdsson, T. Silagadze, A. B. Smit, H. Stefansson, S. Steinberg, J. Suvisaari, S. Tosato, M. Verhage, J. T. Walters, Consortium Multicenter Genetic Studies of Schizophrenia D. F. Levinson, P. V. Gejman, K. S. Kendler, C. Laurent, B. J. Mowry, M. C. O’Donovan, M. J. Owen, A. E. Pulver, B. P. Riley, S. G. Schwab, D. B. Wildenauer, F. Dudbridge, P. Holmans, J. Shi, M. Albus, M. Alexander, D. Campion, D. Cohen, D. Dikeos, J. Duan, P. Eichhammer, S. Godard, M. Hansen, F. B. Lerer, K. Y. Liang, W. Maier, J. Mallet, D. A. Nertney, G. Nestadt, N. Norton, F. A. O’Neill, G. N. Papadimitriou, R. Ribble, A. R. Sanders, J. M. Silverman, D. Walsh, N. M. Williams, B. Wormley, Consortium Psychosis Endophenotypes International, M. J. Arranz, S. Bakker, S. Bender, E. Bramon, D. Collier, B. Crespo-Facorro, J. Hall, C. Iyegbe, A. Jablensky, R. S. Kahn, L. Kalaydjieva, S. Lawrie, C. M. Lewis, K. Lin, D. H. Linszen, I. Mata, A. McIntosh, R. M. Murray, R. A. Ophoff, J. Powell, D. Rujescu, J. Van Os, M. Walshe, M. Weisbrod, D. Wiersma, Consortium Wellcome Trust Case Control, P. Donnelly, I. Barroso, J. M. Blackwell, E. Bramon, M. A. Brown, J. P. Casas, A. P. Corvin, P. Deloukas, A. Duncanson, J. Jankowski, H. S. Markus, C. G. Mathew, C. N. Palmer, R. Plomin, A. Rautanen, S. J. Sawcer, R. C. Trembath, A. C. Viswanathan, N. W. Wood, C. C. Spencer, G. Band, C. Bellenguez, C. Freeman, G. Hellenthal, E. Giannoulatou, M. Pirinen, R. D. Pearson, A. Strange, Z. Su, D. Vukcevic, P. Donnelly, C. Langford, S. E. Hunt, S. Edkins, R. Gwilliam, H. Blackburn, S. J. Bumpstead, S. Dronov, M. Gillman, E. Gray, N. Hammond, A. Jayakumar, O. T. McCann, J. Liddle, S. C. Potter, R. Ravindrarajah, M. Ricketts, A. Tashakkori-Ghanbaria, M. J. Waller, P. Weston, S. Widaa, P. Whittaker, I. Barroso, P. Deloukas, C. G. Mathew, J. M. Blackwell, M. A. Brown, A. P. Corvin, M. I. McCarthy, C. C. Spencer, E. Bramon, A. P. Corvin, M. C. O’Donovan, K. Stefansson, E. Scolnick, S. Purcell, S. A. McCarroll, P. Sklar, C. M. Hultman, and P. F. Sullivan. 2013. ‘Genome-wide association analysis identifies 13 new risk loci for schizophrenia’, Nat Genet, 45: 1150–9.

Robicsek, O., R. Karry, I. Petit, N. Salman-Kesner, F. J. Müller, E. Klein, D. Aberdam, and D. Ben-Shachar. 2013. ‘Abnormal neuronal differentiation and mitochondrial dysfunction in hair follicle-derived induced pluripotent stem cells of schizophrenia patients’, Mol Psychiatry, 18: 1067–76.

Robicsek, Odile, Hila M Ene, Rachel Karry, Ofer Ytzhaki, Eyal Asor, Donna McPhie, Bruce M Cohen, Rotem Ben-Yehuda, Ina Weiner, and Dorit Ben-Shachar. 2017. ‘Isolated Mitochondria Transfer Improves Neuronal Differentiation of Schizophrenia-Derived Induced Pluripotent Stem Cells and Rescues Deficits in a Rat Model of the Disorder’, Schizophr Bull, 44: 432–42.

Rosenfeld, Marina, Hanit Brenner-Lavie, Shunit Gal-Ben Ari, Alexandra Kavushansky, and Dorit Ben-Shachar. 2011. ‘Perturbation in Mitochondrial Network Dynamics and in Complex I Dependent Cellular Respiration in Schizophrenia’, Biological Psychiatry, 69: 980–88.

Rowe, R. Grant, and George Q. Daley. 2019. ‘Induced pluripotent stem cells in disease modelling and drug discovery’, Nature Reviews Genetics, 20: 377–88.

Salta, Evgenia, and Bart De Strooper. 2012. ‘Non-coding RNAs with essential roles in neurodegenerative disorders’, The Lancet Neurology, 11: 189–200.

Sarkar, Anindita, Arianna Mei, Apua C. M. Paquola, Shani Stern, Cedric Bardy, Jason R. Klug, Stacy Kim, Neda Neshat, Hyung Joon Kim, Manching Ku, Maxim N. Shokhirev, David H. Adamowicz, Maria C. Marchetto, Roberto Jappelli, Jennifer A. Erwin, Krishnan Padmanabhan, Matthew Shtrahman, Xin Jin, and Fred H. Gage. 2018. ‘Efficient Generation of CA3 Neurons from Human Pluripotent Stem Cells Enables Modeling of Hippocampal Connectivity In Vitro’, Cell Stem Cell, 22: 684–97.e9.

Sawada, Tomoyo, Thomas E Chater, Yohei Sasagawa, Mika Yoshimura, Noriko Fujimori-Tonou, Kaori Tanaka, Kynon JM Benjamin, Apua Paquola, Jennifer A Erwin, and Yukiko %J Molecular psychiatry Goda. 2020. ’Developmental excitation-inhibition imbalance underlying psychoses revealed by single-cell analyses of discordant twins-derived cerebral organoids’, 25: 2695–711.

Schizophrenia Psychiatric Genome-Wide Association Study, Consortium. 2011a. ‘Genome-wide association study identifies five new schizophrenia loci’, Nature Genetics, 43: 969–76.

Schizophrenia Psychiatric Genome-Wide Association Study, Consortium. 2011a. ‘, Nat Genet, 43: 969–76.

Schizophrenia Working Group of the Psychiatric Genomics, Consortium. 2014. ‘Biological insights from 108 schizophrenia-associated genetic loci’, Nature, 511: 421–27.

Schultz, S. H., S. W. North, and C. G. Shields. 2007. ‘Schizophrenia: a review’, Am Fam Physician, 75: 1821–9.

Sellgren, Carl M., Jessica Gracias, Bradley Watmuff, Jonathan D. Biag, Jessica M. Thanos, Paul B. Whittredge, Ting Fu, Kathleen Worringer, Hannah E. Brown, Jennifer Wang, Ajamete Kaykas, Rakesh Karmacharya, Carleton P. Goold, Steven D. Sheridan, and Roy H. Perlis. 2019. ‘Increased synapse elimination by microglia in schizophrenia patient-derived models of synaptic pruning’, Nature Neuroscience, 22: 374–85.

Singh, Tarjinder, Mitja I. Kurki, David Curtis, Shaun M. Purcell, Lucy Crooks, Jeremy McRae, Jaana Suvisaari, Himanshu Chheda, Douglas Blackwood, Gerome Breen, Olli Pietiläinen, Sebastian S. Gerety, Muhammad Ayub, Moira Blyth, Trevor Cole, David Collier, Eve L. Coomber, Nick Craddock, Mark J. Daly, John Danesh, Marta DiForti, Alison Foster, Nelson B. Freimer, Daniel Geschwind, Mandy Johnstone, Shelagh Joss, Georg Kirov, Jarmo Körkkö, Outi Kuismin, Peter Holmans, Christina M. Hultman, Conrad Iyegbe, Jouko Lönnqvist, Minna Männikkö, Steve A. McCarroll, Peter McGuffin, Andrew M. McIntosh, Andrew McQuillin, Jukka S. Moilanen, Carmel Moore, Robin M. Murray, Ruth Newbury-Ecob, Willem Ouwehand, Tiina Paunio, Elena Prigmore, Elliott Rees, David Roberts, Jennifer Sambrook, Pamela Sklar, David St Clair, Juha Veijola, James T. R. Walters, Hywel Williams, Study Swedish Schizophrenia, Interval Study, D. D. D. Study, Uk K. Consortium, Patrick F. Sullivan, Matthew E. Hurles, Michael C. O’Donovan, Aarno Palotie, Michael J. Owen, and Jeffrey C. Barrett. 2016. ‘Rare loss-of-function variants in SETD1A are associated with schizophrenia and developmental disorders’, Nature Neuroscience, 19: 571–77.

Singh, Tarjinder, Timothy Poterba, David Curtis, Huda Akil, Mariam Al Eissa, Jack D. Barchas, Nicholas Bass, Tim B. Bigdeli, Gerome Breen, Evelyn J. Bromet, Peter F. Buckley, William E. Bunney, Jonas Bybjerg-Grauholm, William F. Byerley, Sinéad B. Chapman, Wei J. Chen, Claire Churchhouse, Nicholas Craddock, Caroline M. Cusick, Lynn DeLisi, Sheila Dodge, Michael A. Escamilla, Saana Eskelinen, Ayman H. Fanous, Stephen V. Faraone, Alessia Fiorentino, Laurent Francioli, Stacey B. Gabriel, Diane Gage, Sarah A. Gagliano Taliun, Andrea Ganna, Giulio Genovese, David C. Glahn, Jakob Grove, Mei-Hua Hall, Eija Hämäläinen, Henrike O. Heyne, Matti Holi, David M. Hougaard, Daniel P. Howrigan, Hailiang Huang, Hai-Gwo Hwu, René S. Kahn, Hyun Min Kang, Konrad J. Karczewski, George Kirov, James A. Knowles, Francis S. Lee, Douglas S. Lehrer, Francesco Lescai, Dolores Malaspina, Stephen R. Marder, Steven A. McCarroll, Andrew M. McIntosh, Helena Medeiros, Lili Milani, Christopher P. Morley, Derek W. Morris, Preben Bo Mortensen, Richard M. Myers, Merete Nordentoft, Niamh L. O’Brien, Ana Maria Olivares, Dost Ongur, Willem H. Ouwehand, Duncan S. Palmer, Tiina Paunio, Digby Quested, Mark H. Rapaport, Elliott Rees, Brandi Rollins, F. Kyle Satterstrom, Alan Schatzberg, Edward Scolnick, Laura J. Scott, Sally I. Sharp, Pamela Sklar, Jordan W. Smoller, Janet L. Sobell, Matthew Solomonson, Eli A. Stahl, Christine R. Stevens, Jaana Suvisaari, Grace Tiao, Stanley J. Watson, Nicholas A. Watts, Douglas H. Blackwood, Anders D. Børglum, Bruce M. Cohen, Aiden P. Corvin, Tõnu Esko, Nelson B. Freimer, Stephen J. Glatt, Christina M. Hultman, Andrew McQuillin, Aarno Palotie, Carlos N. Pato, Michele T. Pato, Ann E. Pulver, David St. Clair, Ming T. Tsuang, Marquis P. Vawter, James T. Walters, Thomas M. Werge, Roel A. Ophoff, Patrick F. Sullivan, Michael J. Owen, Michael Boehnke, Michael C. O’Donovan, Benjamin M. Neale, and Mark J. Daly. 2022. ‘Rare coding variants in ten genes confer substantial risk for schizophrenia’, Nature, 604: 509–16.

Soliman, M. A., F. Aboharb, N. Zeltner, and L. Studer. 2017. ‘Pluripotent stem cells in neuropsychiatric disorders’, Mol Psychiatry, 22: 1241–49.

Srikanth, Priya, Karam Han, Dana G Callahan, Eugenia Makovkina, Christina R Muratore, Matthew A Lalli, Honglin Zhou, Justin D Boyd, Kenneth S Kosik, Dennis J Selkoe, and Tracy L Young-Pearse. 2015. ‘Genomic DISC1 Disruption in hiPSCs Alters Wnt Signaling and Neural Cell Fate’, Cell Reports, 12: 1414–29.

Srikanth, Priya, Valentina N Lagomarsino, Christina R Muratore, Steven C Ryu, Amy He, Walter M Taylor, Constance Zhou, Marlise Arellano, and Tracy L %J Translational psychiatry Young-Pearse. 2018. ’Shared effects of DISC1 disruption and elevated WNT signaling in human cerebral organoids’, 8: 1–14.

Stachowiak, EK, CA Benson, ST Narla, A Dimitri, LE Chuye, S Dhiman, K Harikrishnan, S Elahi, D Freedman, and KJ %J Translational psychiatry Brennand. 2017. ’Cerebral organoids reveal early cortical maldevelopment in schizophrenia—computational anatomy and genomics, role of FGFR1’, 7: 1–24.

Stern, Shani, Shong Lau, Andreea Manole, Idan Rosh, Menahem Percia, Ran Ben Ezer, Maxim N. Shokhirev, Fan Qiu, Simon Schafer, Abed Mansour, Tchelet Stern, Pola Ofer, Yam Stern, Ana Mendes Diniz, Lynne Randolph Moore, Ritu Nayak, Aidan Aicher, Amanda Rhee, Thomas L. Wong, Thao Nguyen, Sara B. Linker, Beate Winner, Beatriz C. Freitas, Eugenia Jones, Cedric Bardy, Alexis Brice, Juergen Winkler, Maria C. Marchetto, and Fred H. Gage. 2022. ’Reduced synaptic activity and dysregulated extracellular matrix pathways are common phenotypes in midbrain neurons derived from sporadic and mutation-associated Parkinson’s disease patients’: 2021.12.31.474654.

Stern, Shani, Sara Linker, Krishna C. Vadodaria, Maria C. Marchetto, and Fred H. Gage. 2018. ’Prediction of response to drug therapy in psychiatric disorders’, 8: 180031.

Stern, Shani, Anindita Sarkar, Dekel Galor, Tchelet Stern, Arianna Mei, Yam Stern, Ana P. D. Mendes, Lynne Randolph-Moore, Guy Rouleau, Anne G. Bang, Renata Santos, Martin Alda, Maria C. Marchetto, and Fred H. Gage. 2020. ‘A Physiological Instability Displayed in Hippocampal Neurons Derived From Lithium-Nonresponsive Bipolar Disorder Patients’, Biological Psychiatry, 88: 150–58.

Stern, Shani, Anindita Sarkar, Tchelet Stern, Arianna Mei, Ana P. D. Mendes, Yam Stern, Gabriela Goldberg, Dekel Galor, Thao Nguyen, Lynne Randolph-Moore, Yongsung Kim, Guy Rouleau, Anne Bang, Martin Alda, Renata Santos, Maria C. Marchetto, and Fred H. Gage. 2020. ‘Mechanisms Underlying the Hyperexcitability of CA3 and Dentate Gyrus Hippocampal Neurons Derived From Patients With Bipolar Disorder’, Biological Psychiatry, 88: 139–49.

Stern, Shani, Lei Zhang, Meiyan Wang, Rebecca Wright, Diogo Cordeiro, David Peles, Yuqing Hang, Ana P. D. Mendes, Tithi Baul, Julien Roth, Shashank Coorapati, Marco Boks, Hilleke Hulshoff Pol, Kristen J. Brennand, Janos M Réthelyi, René S. Kahn, Maria C. Marchetto, and Fred H. Gage. 2022. ’Monozygotic twins discordant for schizophrenia differ in maturation and synaptic transmission’: 2022.05.13.491776.

Stilo, S. A., and R. M. Murray. 2010. ‘The epidemiology of schizophrenia: replacing dogma with knowledge’, Dialogues Clin Neurosci, 12: 305–15.

Sullivan, Patrick F., Mark J. Daly, and Michael O’Donovan. 2012. ‘Genetic architectures of psychiatric disorders: the emerging picture and its implications’, Nature Reviews Genetics, 13: 537–51.

Sullivan, Patrick F., Kenneth S. Kendler, and Michael C. Neale. 2003. ‘Schizophrenia as a Complex Trait: Evidence From a Meta-analysis of Twin Studies’, Arch Gen Psychiatry, 60: 1187–92.

Szabo, Attila, Ibrahim A. Akkouh, Matthieu Vandenberghe, Jordi Requena Osete, Timothy Hughes, Vivi Heine, Olav B. Smeland, Joel C. Glover, Ole A. Andreassen, and Srdjan Djurovic. 2021. ‘A human iPSC-astroglia neurodevelopmental model reveals divergent transcriptomic patterns in schizophrenia’, Transl Psychiatry, 11: 554.

Takahashi, K., and S. Yamanaka. 2006. ‘Induction of pluripotent stem cells from mouse embryonic and adult fibroblast cultures by defined factors’, Cell, 126: 663–76.

Tandon, R., W. Gaebel, D. M. Barch, J. Bustillo, R. E. Gur, S. Heckers, D. Malaspina, M. J. Owen, S. Schultz, M. Tsuang, J. Van Os, and W. Carpenter. 2013. ‘Definition and description of schizophrenia in the DSM-5’, Schizophr Res, 150: 3–10.

Tandon, R., H. A. Nasrallah, and M. S. Keshavan. 2010. ‘Schizophrenia, “just the facts” 5. Treatment and prevention. Past, present, and future’, Schizophr Res, 122: 1–23.

Tian, T., Z. Wei, X. Chang, Y. Liu, R. E. Gur, P. M. A. Sleiman, and H. Hakonarson. 2018. ‘The Long Noncoding RNA Landscape in Amygdala Tissues from Schizophrenia Patients’, EBioMedicine, 34: 171–81.

Tiihonen, J, M Koskuvi, M Storvik, I Hyötyläinen, Y Gao, KA Puttonen, R Giniatullina, E Poguzhelskaya, I Ojansuu, and O Vaurio. 2019. ”Sex-specific transcriptional and proteomic signatures in schizophrenia. Nat Commun 10 (1): 3933.” In.

Topol, Aaron, Shijia Zhu, Ngoc Tran, Anthony Simone, Gang Fang, and Kristen J. Brennand. 2015. ‘Altered WNT Signaling in Human Induced Pluripotent Stem Cell Neural Progenitor Cells Derived from Four Schizophrenia Patients’, Biological Psychiatry, 78: e29–e34.

Toyoshima, M., W. Akamatsu, Y. Okada, T. Ohnishi, S. Balan, Y. Hisano, Y. Iwayama, T. Toyota, T. Matsumoto, N. Itasaka, S. Sugiyama, M. Tanaka, M. Yano, B. Dean, H. Okano, and T. Yoshikawa. 2016. ‘Analysis of induced pluripotent stem cells carrying 22q11.2 deletion’, Transl Psychiatry, 6: e934–e34.

Trifu, Simona Corina, Bianca Kohn, Andrei Vlasie, and Bogdan-Eduard Patrichi. 2020. ‘Genetics of schizophrenia (Review)’, Experimental and therapeutic medicine, 20: 3462–68.

Trubetskoy, Vassily, Antonio F. Pardiñas, Ting Qi, Georgia Panagiotaropoulou, Swapnil Awasthi, Tim B. Bigdeli, Julien Bryois, Chia-Yen Chen, Charlotte A. Dennison, Lynsey S. Hall, Max Lam, Kyoko Watanabe, Oleksandr Frei, Tian Ge, Janet C. Harwood, Frank Koopmans, Sigurdur Magnusson, Alexander L. Richards, Julia Sidorenko, Yang Wu, Jian Zeng, Jakob Grove, Minsoo Kim, Zhiqiang Li, Georgios Voloudakis, Wen Zhang, Mark Adams, Ingrid Agartz, Elizabeth G. Atkinson, Esben Agerbo, Mariam Al Eissa, Margot Albus, Madeline Alexander, Behrooz Z. Alizadeh, Köksal Alptekin, Thomas D. Als, Farooq Amin, Volker Arolt, Manuel Arrojo, Lavinia Athanasiu, Maria Helena Azevedo, Silviu A. Bacanu, Nicholas J. Bass, Martin Begemann, Richard A. Belliveau, Judit Bene, Beben Benyamin, Sarah E. Bergen, Giuseppe Blasi, Julio Bobes, Stefano Bonassi, Alice Braun, Rodrigo Affonseca Bressan, Evelyn J. Bromet, Richard Bruggeman, Peter F. Buckley, Randy L. Buckner, Jonas Bybjerg-Grauholm, Wiepke Cahn, Murray J. Cairns, Monica E. Calkins, Vaughan J. Carr, David Castle, Stanley V. Catts, Kimberley D. Chambert, Raymond C. K. Chan, Boris Chaumette, Wei Cheng, Eric F. C. Cheung, Siow Ann Chong, David Cohen, Angèle Consoli, Quirino Cordeiro, Javier Costas, Charles Curtis, Michael Davidson, Kenneth L. Davis, Lieuwe de Haan, Franziska Degenhardt, Lynn E. DeLisi, Ditte Demontis, Faith Dickerson, Dimitris Dikeos, Timothy Dinan, Srdjan Djurovic, Jubao Duan, Giuseppe Ducci, Frank Dudbridge, Johan G. Eriksson, Lourdes Fañanás, Stephen V. Faraone, Alessia Fiorentino, Andreas Forstner, Josef Frank, Nelson B. Freimer, Menachem Fromer, Alessandra Frustaci, Ary Gadelha, Giulio Genovese, Elliot S. Gershon, Marianna Giannitelli, Ina Giegling, Paola Giusti-Rodríguez, Stephanie Godard, Jacqueline I. Goldstein, Javier González Peñas, Ana González-Pinto, Srihari Gopal, Jacob Gratten, Michael F. Green, Tiffany A. Greenwood, Olivier Guillin, Sinan Gülöksüz, Raquel E. Gur, Ruben C. Gur, Blanca Gutiérrez, Eric Hahn, Hakon Hakonarson, Vahram Haroutunian, Annette M. Hartmann, Carol Harvey, Caroline Hayward, Frans A. Henskens, Stefan Herms, Per Hoffmann, Daniel P. Howrigan, Masashi Ikeda, Conrad Iyegbe, Inge Joa, Antonio Julià, Anna K. Kähler, Tony Kam-Thong, Yoichiro Kamatani, Sena Karachanak-Yankova, Oussama Kebir, Matthew C. Keller, Brian J. Kelly, Andrey Khrunin, Sung-Wan Kim, Janis Klovins, Nikolay Kondratiev, Bettina Konte, Julia Kraft, Michiaki Kubo, Vaidutis Kučinskas, Zita Ausrele Kučinskiene, Agung Kusumawardhani, Hana Kuzelova-Ptackova, Stefano Landi, Laura C. Lazzeroni, Phil H. Lee, Sophie E. Legge, Douglas S. Lehrer, Rebecca Lencer, Bernard Lerer, Miaoxin Li, Jeffrey Lieberman, Gregory A. Light, Svetlana Limborska, Chih-Min Liu, Jouko Lönnqvist, Carmel M. Loughland, Jan Lubinski, Jurjen J. Luykx, Amy Lynham, Milan Macek, Andrew Mackinnon, Patrik K. E. Magnusson, Brion S. Maher, Wolfgang Maier, Dolores Malaspina, Jacques Mallet, Stephen R. Marder, Sara Marsal, Alicia R. Martin, Lourdes Martorell, Manuel Mattheisen, Robert W. McCarley, Colm McDonald, John J. McGrath, Helena Medeiros, Sandra Meier, Bela Melegh, Ingrid Melle, Raquelle I. Mesholam-Gately, Andres Metspalu, Patricia T. Michie, Lili Milani, Vihra Milanova, Marina Mitjans, Espen Molden, Esther Molina, María Dolores Molto, Valeria Mondelli, Carmen Moreno, Christopher P. Morley, Gerard Muntané, Kieran C. Murphy, Inez Myin-Germeys, Igor Nenadić, Gerald Nestadt, Liene Nikitina-Zake, Cristiano Noto, Keith H. Nuechterlein, Niamh Louise O’Brien, F. Anthony O’Neill, Sang-Yun Oh, Ann Olincy, Vanessa Kiyomi Ota, Christos Pantelis, George N. Papadimitriou, Mara Parellada, Tiina Paunio, Renata Pellegrino, Sathish Periyasamy, Diana O. Perkins, Bruno Pfuhlmann, Olli Pietiläinen, Jonathan Pimm, David Porteous, John Powell, Diego Quattrone, Digby Quested, Allen D. Radant, Antonio Rampino, Mark H. Rapaport, Anna Rautanen, Abraham Reichenberg, Cheryl Roe, Joshua L. Roffman, Julian Roth, Matthias Rothermundt, Bart P. F. Rutten, Safaa Saker-Delye, Veikko Salomaa, Julio Sanjuan, Marcos Leite Santoro, Adam Savitz, Ulrich Schall, Rodney J. Scott, Larry J. Seidman, Sally Isabel Sharp, Jianxin Shi, Larry J. Siever, Engilbert Sigurdsson, Kang Sim, Nora Skarabis, Petr Slominsky, Hon-Cheong So, Janet L. Sobell, Erik Söderman, Helen J. Stain, Nils Eiel Steen, Agnes A. Steixner-Kumar, Elisabeth Stögmann, William S. Stone, Richard E. Straub, Fabian Streit, Eric Strengman, T. Scott Stroup, Mythily Subramaniam, Catherine A. Sugar, Jaana Suvisaari, Dragan M. Svrakic, Neal R. Swerdlow, Jin P. Szatkiewicz, Thi Minh Tam Ta, Atsushi Takahashi, Chikashi Terao, Florence Thibaut, Draga Toncheva, Paul A. Tooney, Silvia Torretta, Sarah Tosato, Gian Battista Tura, Bruce I. Turetsky, Alp Üçok, Arne Vaaler, Therese van Amelsvoort, Ruud van Winkel, Juha Veijola, John Waddington, Henrik Walter, Anna Waterreus, Bradley T. Webb, Mark Weiser, Nigel M. Williams, Stephanie H. Witt, Brandon K. Wormley, Jing Qin Wu, Zhida Xu, Robert Yolken, Clement C. Zai, Wei Zhou, Feng Zhu, Fritz Zimprich, Eşref Cem Atbaşoğlu, Muhammad Ayub, Christian Benner, and Alessandro Bertolino. 2022. ‘Mapping genomic loci implicates genes and synaptic biology in schizophrenia’, Nature, 604: 502–08.

Uffelmann, Emil, Qin Qin Huang, Nchangwi Syntia Munung, Jantina de Vries, Yukinori Okada, Alicia R. Martin, Hilary C. Martin, Tuuli Lappalainen, and Danielle Posthuma. 2021. ‘Genome-wide association studies’, Nature Reviews Methods Primers, 1: 59.

van Dongen, J., and D. I. Boomsma. 2013. ‘The evolutionary paradox and the missing heritability of schizophrenia’, Am J Med Genet B Neuropsychiatr Genet, 162b: 122–36.

van Haren, N. E., H. G. Schnack, W. Cahn, M. P. van den Heuvel, C. Lepage, L. Collins, A. C. Evans, H. E. Hulshoff Pol, and R. S. Kahn. 2011. ‘Changes in cortical thickness during the course of illness in schizophrenia’, Arch Gen Psychiatry, 68: 871–80.

Warre-Cornish, Katherine, Leo Perfect, Roland Nagy, Rodrigo RR Duarte, Matthew J Reid, Pooja Raval, Annett Mueller, Amanda L Evans, Amalie Couch, and Cédric %J Science advances Ghevaert. 2020. ’Interferon-γ signaling in human iPSC–derived neurons recapitulates neurodevelopmental disorder phenotypes’, 6: eaay9506.

Wen, Zhexing, Ha Nam Nguyen, Ziyuan Guo, Matthew A. Lalli, Xinyuan Wang, Yijing Su, Nam-Shik Kim, Ki-Jun Yoon, Jaehoon Shin, Ce Zhang, Georgia Makri, David Nauen, Huimei Yu, Elmer Guzman, Cheng-Hsuan Chiang, Nadine Yoritomo, Kozo Kaibuchi, Jizhong Zou, Kimberly M. Christian, Linzhao Cheng, Christopher A. Ross, Russell L. Margolis, Gong Chen, Kenneth S. Kosik, Hongjun Song, and Guo-li Ming. 2014. ‘Synaptic dysregulation in a human iPS cell model of mental disorders’, Nature, 515: 414–18.

Wildgust, H. J., R. Hodgson, and M. Beary. 2010. ‘The paradox of premature mortality in schizophrenia: new research questions’, J Psychopharmacol, 24: 9–15.

Windrem, Martha S., Mikhail Osipovitch, Zhengshan Liu, Janna Bates, Devin Chandler-Militello, Lisa Zou, Jared Munir, Steven Schanz, Katherine McCoy, Robert H. Miller, Su Wang, Maiken Nedergaard, Robert L. Findling, Paul J. Tesar, and Steven A. Goldman. 2017. ‘Human iPSC Glial Mouse Chimeras Reveal Glial Contributions to Schizophrenia’, Cell Stem Cell, 21: 195–208.e6.

Wrobel, Carolyn N, Christopher A Mutch, Sruthi Swaminathan, Makoto M Taketo, and Anjen %J Developmental biology Chenn. 2007. ’Persistent expression of stabilized β-catenin delays maturation of radial glial cells into intermediate progenitors’, 309: 285–97.

Xie, Yan, Evan Xu, and Ziyad Al-Aly. 2022. ’Risks of mental health outcomes in people with covid-19: cohort study’, 376: e068993.

Yoon, Ki-Jun, Ha Nam Nguyen, Gianluca Ursini, Fengyu Zhang, Nam-Shik Kim, Zhexing Wen, Georgia Makri, David Nauen, Joo Heon Shin, Youngbin Park, Raeeun Chung, Eva Pekle, Ce Zhang, Maxwell Towe, Syed Mohammed Qasim Hussaini, Yohan Lee, Dan Rujescu, David St. Clair, Joel E Kleinman, Thomas M Hyde, Gregory Krauss, Kimberly M Christian, Judith L Rapoport, Daniel R Weinberger, Hongjun Song, and Guo-li Ming. 2014. ‘Modeling a Genetic Risk for Schizophrenia in iPSCs and Mice Reveals Neural Stem Cell Deficits Associated with Adherens Junctions and Polarity’, Cell Stem Cell, 15: 79–91.

Zhang, C., E. Tannous, and J. J. Zheng. 2019. ‘Oxidative stress upregulates Wnt signaling in human retinal microvascular endothelial cells through activation of disheveled’, J Cell Biochem, 120: 14044–54.

Zhang, Xiaojuan, Lijun Pei, Runting Li, Wei Zhang, Hua Yang, Yongchao Li, Yu Guo, Pingping Tan, Jingdong J. Han, Xiaoying Zheng, and Runlin Z. Ma. 2015. ‘Spina bifida in fetus is associated with an altered pattern of DNA methylation in placenta’, Journal of Human Genetics, 60: 605–11.

Zhao, Jian, Baoshan Cai, Zhugui Shao, Lei Zhang, Yi Zheng, Chunhong Ma, Fan Yi, Bingyu Liu, and Chengjiang Gao. 2021. ‘TRIM26 positively regulates the inflammatory immune response through K11-linked ubiquitination of TAB1’, Cell Death & Differentiation, 28: 3077–91.

Zheutlin, A. B., J. Dennis, R. Karlsson Linnér, A. Moscati, N. Restrepo, P. Straub, D. Ruderfer, V. M. Castro, C. Y. Chen, T. Ge, L. M. Huckins, A. Charney, H. L. Kirchner, E. A. Stahl, C. F. Chabris, L. K. Davis, and J. W. Smoller. 2019. ‘Penetrance and Pleiotropy of Polygenic Risk Scores for Schizophrenia in 106,160 Patients Across Four Health Care Systems’, Am J Psychiatry, 176: 846–55.

Zuk, Or, Eliana Hechter, Shamil R. Sunyaev, and Eric S. Lander. 2012. ’The mystery of missing heritability: Genetic interactions create phantom heritability’, 109: 1193–98.

